# BAHD news from *Euphorbia peplus*: identification of acyltransferase enzymes involved in ingenane diterpenoid biosynthesis

**DOI:** 10.1101/2025.03.03.641262

**Authors:** Carsten Schotte, Matilde Florean, Tomasz Czechowski, Alison Gilday, Ryan M. Alam, Kerstin Ploss, Jens Wurlitzer, Yi Li, Prashant Sonawane, Ian A. Graham, Sarah E. O’Connor

## Abstract

- The plant family *Euphorbiaceae* are an abundant source of structurally complex diterpenoids, many of which have reported anti-cancer, anti-HIV, and anti-inflammatory activities. Among these, ingenol-3-angelate (**1a**; tradename: Picato^®^), isolated from *Euphorbia peplus*, has potent anti-tumour activity.
- Here we report the discovery and characterization of the first genes linked to the committed steps of ingenol-3-angelate (**1a**) biosynthesis in *Euphorbia peplus*. We identified two genes, the products of which catalyse the addition of angelyl-CoA (**9a**) to the ingenol (**5**) scaffold to produce ingenol-3-angelate (**1a**).
- We demonstrate using VIGS that just one of these genes, *EpBAHD-08*, is essential for this angeloylation in *E. peplus*. VIGS of the second gene, *EpBAHD-06*, has a significant effect on jatrophanes rather than ingenanes in *E. peplus*.
- We also identified three genes whose products can catalyse acetylation of ingenol-3-angelate (**1a**) to ingenol-3-angelate-20-acetate (**2**). In this case VIGS indicates considerable functional redundancy in the *E. peplus* genome of genes encoding this enzymatic step.
- This work paves the way for increasing ingenol-3-angelate (**1a**) levels *in planta* and provides a foundation for the discovery of the remaining genes in the biosynthetic pathway of these important molecules.

## Introduction

The *Euphorbiaceae* are one of the largest families of plants with more than 7500 species reported to date (1, 2). Notably, the majority of *Euphorbiaceae* plants produce a milky latex that is rich in biologically active diterpenoids (1). From the genus *Euphorbia*, >1500 diterpenoids with more than 30 different skeletal backbones have been isolated (2–4). These skeletal backbones have exceptional structural complexity, and members of this natural product family range in the number of ring systems, degree of oxygenation, stereochemistry, and esterification pattern (5).

Clinically important diterpenoids from the *Euphorbia* genus include resiniferatoxin (**3**), a transient receptor vanilloid 1 (TRPV1) agonist, that is currently being investigated for treatment of an overactive bladder and chronic pain (phase III clinical trial), and prostratin (**4**), which may have applications in clearing latent virus reservoirs in HIV infections (Figure 1A) (2, 6, 7). Ingenol-3-angelate (**1a**) is perhaps the most well-known diterpene that is produced by *Euphorbia* (*Euphorbia peplus*; Figure 1B). This compound was approved in 2012 by the FDA for the treatment of the precancerous skin condition actinic keratosis (Picato^®^) but discontinued in 2020 in the course of a phase IV clinical trial when it appeared that this compound increased the incidence of skin cancer (8). However, ingenol-3-angelate (**1a**) is still being explored in the treatment of HIV infections (9). Additionally, members of this type of diterpenoid (ingenane class) appear to be a rich source of biologically active compounds (8). Therefore, ingenol-3-angelate (**1a**) and related compounds continue to be of high interest for pharmaceutical development.

**Figure 1.**
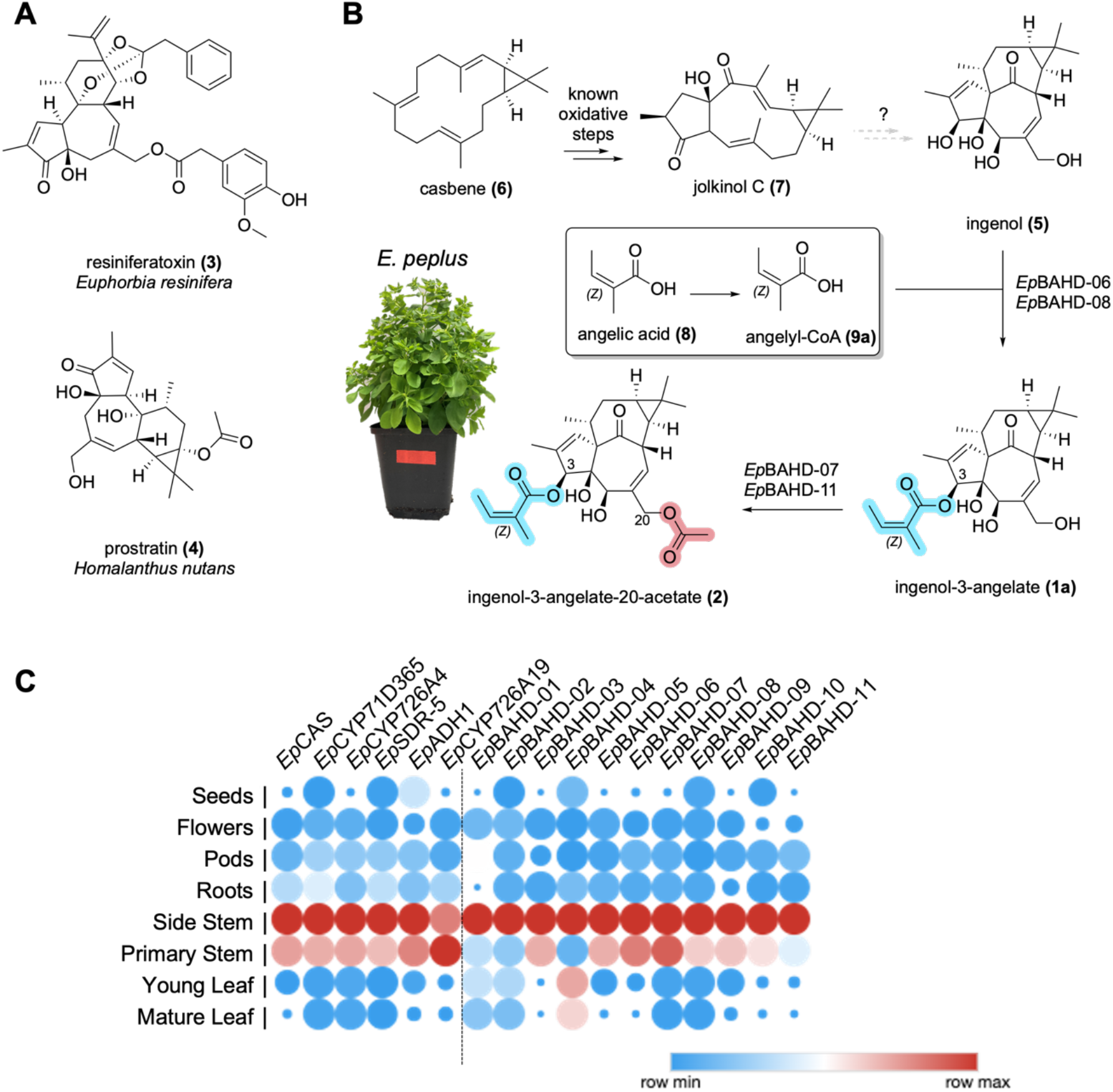
Diterpene biosynthesis in the *Euphorbia* genus. **A)** Clinically important diterpenoids isolated from various *Euphorbia* species. **B)** Known, proposed and newly discovered steps in *E. peplus* diterpene biosynthesis. Genes identified in this study catalyse formation of ingenol-3-angelate (**1a**) from ingenol (**5**) (*Ep*BAHD-06 and *Ep*BAHD-08); and ingenol-3-angelate-20-acetate (**2**) from ingenol-3-angelate (**1a**) (*Ep*BAHD-07 and *Ep*BAHD-11). **C)** Expression profiles of genes involved in jolkinol C (**7**) biosynthesis and BAHD-acyltransferases identified in the course of this study. Red corresponds to maximum expression and blue corresponds to minimum expression levels. Dot size in dependence of relative expression levels across the seven tissues, measured in fragments per kilobase of exon per million of mapped fragments (FPKM).

*Euphorbia* derived diterpenoids typically accumulate to low levels *in planta*: ingenol-3-angelate (**1a**) is present at 1 mg/kg in aerial tissues (10). Chemical syntheses of these structurally complex diterpenes have been reported, but these methods suffer from low yields and long linear reaction sequences (> 10 steps) (11, 12). While semisynthetic approaches towards **1a** from more readily available plant intermediates have been reported (*e*.*g*., 3 steps from ingenol (**5**); Supplementary Figure S1), these approaches still depend on expensive and non-abundant starting materials (13).

Therefore, new methods are needed to produce complex diterpenoids at scale. The use of metabolic engineering methods to produce these compounds in microbial hosts such as yeast could potentially meet this need. Efforts to engineer *Euphorbia* diterpenoid production, however, are limited since the biosynthesis of these terpenes is not well-understood, and most biosynthetic genes that are responsible for the production of these compounds have not been identified.

It is believed that all *Euphorbia*-specific diterpenoids derive from the bicyclic diterpene casbene (**6**) (Figure 1B), which is the initial product of the class I diterpene synthase, casbene synthase (*Ep*CAS) (14). The casbene-derived *Euphorbia* diterpenoids are classified by increasing cyclization of the diterpene backbone, giving rise to jatrophane, lathyrane, tigliane, daphnane, and ingenane classes, with the latter being the most complex diterpenoids (2, 5) (Supplementary Figure S2). Only the biosynthetic genes that convert casbene (**6**) to the simple lathyrane type diterpenoids jolkinol C (**7**) (Figure 1B; Supplementary Figure S3) and jolkinol E (**S3**) have recently been identified (Supplementary Figure S3) (5, 15–17). Silencing of some of these genes in *E. peplus* using virus induced gene silencing (VIGS) confirmed the role of casbene as precursor and suggested that jolkinol C (**7**) may be an on pathway intermediate for ingenol-3-angelate (**1a**), but the majority of downstream acting biosynthetic genes in this pathway and related diterpenoid pathways are unknown (18). Notably, production of jolkinol C (**7**) in baker’s yeast at titres of 800 µg/mL has been reported, suggesting that production by heterologous reconstitution for this class of diterpenes is feasible (17), if subsequent biosynthetic steps are elucidated.

Here, we identify five acyltransferases that derivatize the ingenol (**5**) scaffold to form both ingenol-3-angelate (**1a**) and ingenol-3-angelate-20-acetate (**2**) (Figure 1B), the last predicted enzymatic steps in this biosynthetic network. Transient gene expression in the heterologous host plant *Nicotiana benthamiana* and *in vitro* enzyme assays confirm catalytic activity towards ingenane production. Notably, with exception of two acyltransferases from *Euphorbia lathyris* (26), no acyltransferases had previously been identified in any *Euphorbia* diterpene pathways despite the fact that acylation is one of the most predominant decorations of the *Euphorbia* diterpenoids (Supplementary Figure S4). This discovery fills a gap in the biosynthesis of this important class of compounds, and furthermore, sets the stage for further mining of omics data to identify the remaining missing genes involved in ingenol-3-angelate (**1a**) biosynthesis.

## Results and discussion

*E. peplus* was cultivated in climate chambers until mature seeds ripened. Eight weeks after initiating cultivation, identical tissue samples were collected for both metabolomics and transcriptomics (mature leaves, young leaves, primary stem, side stem, pods, flowers, roots, seeds [mature seeds were collected later after ripening], and latex). Metabolomics analysis revealed the presence of ingenol-3-angelate (**1a**) in all tested tissues, but **1a** was predominantly found in the latex (Supplementary Figure S5). Note, that no RNA could be isolated from latex over the course of this study, preventing transcriptomic analysis of this material. A similar analysis for **2** indicated that this metabolite was also predominantly located in the latex, but again, traces of **2** were found in all tissues (Supplementary Figure S6).

### Gene candidate identification and expression in Nicotiana benthamiana

An analysis of bulk tissue transcriptomic data revealed that the jolkinol E (**S3**) biosynthetic genes are primarily expressed in the primary stem and side stem (Figure 1C). Based on this, we identified BAHD-acyltransferase genes in these tissues that co-express with casbene synthase (Supplementary Table S1) for further functional analysis in tobacco and *in vitro*.

A total of 11 BAHD-acyltransferase candidate genes were selected for functional characterization. Each gene was transiently expressed in *Nicotiana benthamiana* along with an angelyl-CoA ligase gene that had been previously shown to catalyse formation of angelyl-CoA (**9a**) in yeast (*Ep*CCL2) (20). After transfection of the acyl transferase candidate and angelyl-CoA ligase genes in *N. benthamiana*, the substrates ingenol (**5**) and angelic acid (**8**) were infiltrated into the leaf. Leaf tissue was then analyzed by LC-MS to assess the conversion of ingenol (**5**) to ingenol-3-angelate (**1a**). From this assay, we showed that two genes, *Ep*BAHD-06 and *Ep*BAHD-08, catalyse acylation of ingenol (**5**) to form ingenol-3-angelate (**1a**) based on comparison with an authentic standard (Figure 2A and 2B). Intriguingly, both enzymes produced a minor second product that showed identical mass and fragmentation pattern as ingenol-3-angelate (**1a**) (Supplementary Figure S7).

**Figure 2.**
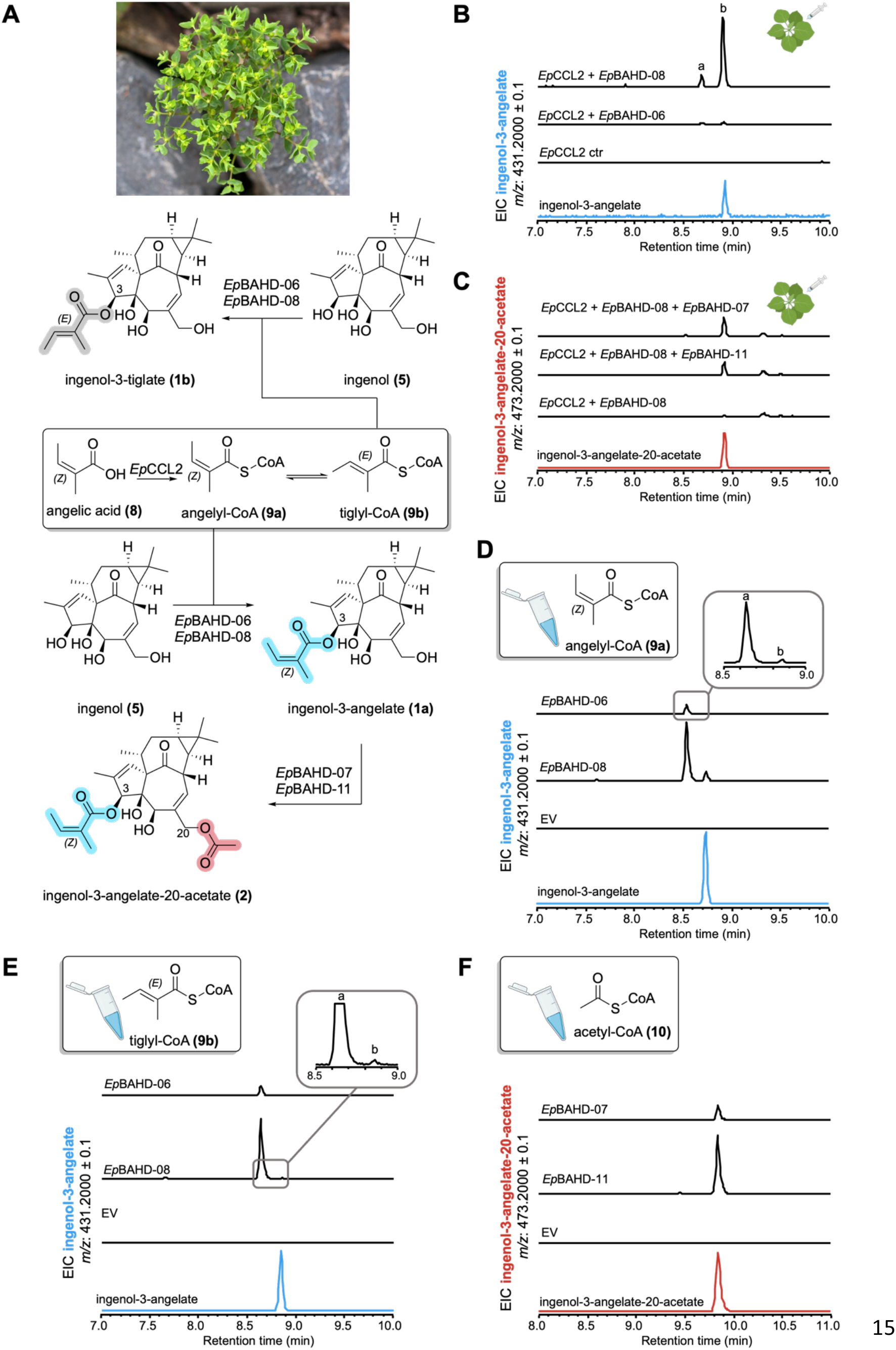
Enzyme activity assays of acyltransferases. **A)** Enzymatic reactions observed in this study. **B)** Tobacco infiltration of a dedicated angelyl-CoA ligase (*Ep*CCL2) (24), angelic acid (**8**), ingenol (**5**) and *Ep*BAHD-06 and *Ep*BAHD-08. Both enzymes catalyse formation of ingenol-3-angelate (**1a**) *in planta* (peak B). **C)** Tobacco infiltration of a dedicated angelyl-CoA ligase (*Ep*CC2L), angelic acid (**8**), ingenol (**5**), *Ep*BAHD-08 (together affording ingenol-3-angelate [**1a**]) and *Ep*BAHD-07 and *Ep*BAHD-11. Both *Ep*BAHD-07 and *Ep*BAHD-11 catalyse formation of ingenol-3-angelate-20-acetate (**2**). Note that LCMS methods in panel B) and C) are different and that **1a** and **2** can be differentiated. **D)** *In vitro* assays with *Ep*BAHD-06 and *Ep*BAHD-08 using angelyl-CoA (**9a**) and ingenol (**5**) as substrate leads to formation of ingenol-3-angelate (**1a**) as a minor product (peak B), with the isomer ingenol-3-tiglate (**1b**) (peak A) as the major product. **E)** *In vitro* assays with *Ep*BAHD-06 and *Ep*BAHD-08 using tiglyl-CoA (**9b**) and ingenol (**5**) as substrate leads to formation of ingenol-3-tiglate (**1b**) (peak A). **F)** *In vitro* assays with *Ep*BAHD-07 and *Ep*BAHD-11 using ingenol-3-angelate (**1a**) and acetyl-CoA (**10**) as substrates leads to formation of ingenol-3-angelate-20-acetate (**2**).

Since angelic acid (**8**) contains an alkene with *Z* configuration, we reasoned that the alkene might readily isomerize to the more stable *E* isomer, tiglic acid (Figure 2A). The *E* to *Z* isomerization of angelic acid (**8**) has already been reported during the synthetic angeloylation of ingenol (**5**) (13). Therefore, we reasoned that the minor product is ingenol-3-tiglate (**1b**) (Figure 2A).

We next assessed whether the angelyl-CoA ligase gene (*Ep*CCL2) is required for production of ingenol-3-angelate (**1a**) in *N. benthamiana*. Interestingly, omission of the angelyl-CoA-ligase had no significant impact on ingenol-3-angelate (**1a**) formation, suggesting that *N. benthamiana* can convert angelic acid (**8**) to angelyl-CoA (**9a**) using endogenous enzyme (Supplementary Figure S8). Notably, addition of angelic acid was absolutely required for formation of **1a**, suggesting that *N. benthamiana* does not have the required pools of this acyl donor (Supplementary Figure S9).

To screen for downstream acyltransferase activity, *Ep*BAHD-08 was again expressed in *N. benthamiana*, but this time together with the other acyltransferase candidates. This assay revealed that in combination with ingenol (**5**) and *Ep*BAHD-08, both *Ep*BAHD-07 and *Ep*BAHD-11 catalyse formation of ingenol-3-angelate-20-acetate (**2**), another well-known, biologically active diterpenoid derivative from *E. peplus* (Alves et al, 2022) (Figure 2C).

### E. coli expression and in vitro assays

*In vitro* enzyme assays using purified recombinant proteins were performed to validate the catalytic activity observed after expression of these genes in *N. benthamiana* leaves. All four BAHD-acyltransferases (*Ep*BAHD-06, *Ep*BAHD-07, *Ep*BAHD-08 and *Ep*BAHD-11) were recombinantly produced in *E. coli*. Angelyl-CoA (**9a**) and tiglyl-CoA (**9b**) were chemically synthesized.

In contrast to assays performed *in planta*, in these *in vitro* assays, incubation of ingenol (**5**) with angelyl-CoA (**9a**) and *Ep*BAHD-06 or *Ep*BAHD-08 led to formation of ingenol-3-angelate (**1a**) as a minor product (Figure 2D). The major product was the previously observed isomer of ingenol-3- angelate (**1a**), which we had tentatively assumed to be ingenol-3-tiglate (**1b**). To confirm this, tiglyl-CoA (**9b**) was also chemically synthesized and then incubated with ingenol (**5**) and *Ep*BAHD-06 or *Ep*BAHD-08. The resulting product was identical to the major product observed in the assay performed with angelyl-CoA (**9a**) (Figure 2E). These *in vitro* assays therefore suggest that the isomerization of the alkene moiety to the more stable *E* isomer likely occurs on angelyl-CoA (**9a**), before transfer of the acyl group to ingenol (**5**).

Finally, incubation of ingenol-3-angelate (**1a**) with acetyl-CoA (**10**) and *Ep*BAHD-07 or *Ep*BAHD-11 led to formation of ingenol-3-angelate-20-acetate (**2**) (Figure 2F).

Next, we performed a phylogenetic analysis of the eleven gene candidates. A multiple sequence alignment was done with 104 functionally annotated acyltransferases from other plants and using two fungal acyltransferase genes as outgroup (Figure 3; Supplementary Figure S10). Plant BAHD acyltransferases group into six distinct clades (21, 22) and the tested acyltransferases from *Euphorbia* form three distinct subgroups within clade 3, a clade generally known to be involved in plant specialized metabolism (22). Notably, *Ep*BAHD-06/*Ep*BAHD-08 and *Ep*BAHD-07/*Ep*BAHD-11 respectively, group in different subclades. Future studies should systematically investigate the functional activity of acyltransferases grouping in other *Euphorbia* specific subclades, to unveil additional acyltransferases involved in the diverse acyltransferase chemistry observed in *Euphorbia* diterpenoids.

**Figure 3.**
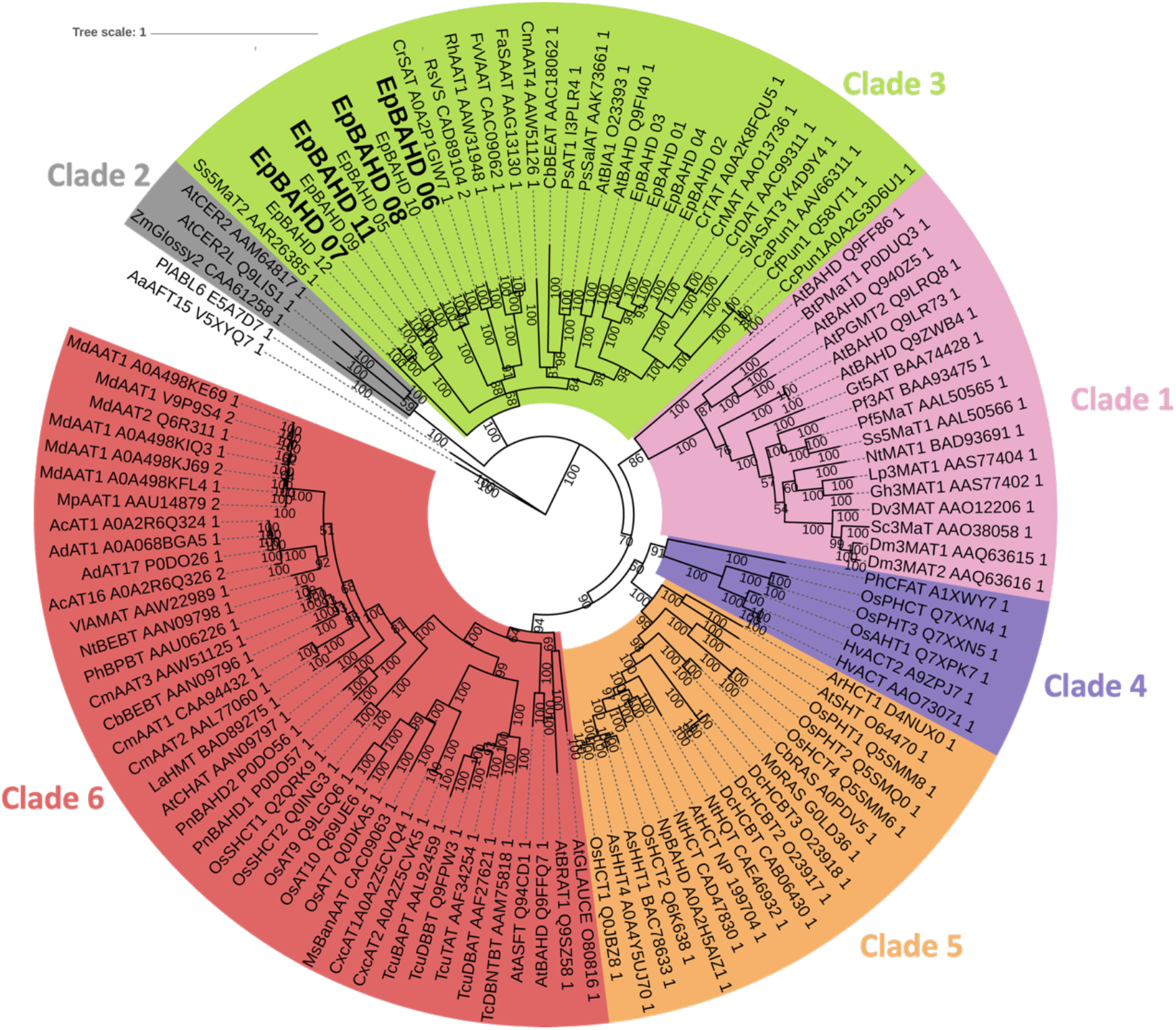
Phylogenetic analysis of acyltransferases identified in this study. Sequence alignment was performed using Muscle v3.8.425 (24). The displayed gene tree was then constructed with Bayesian analyses using MrBayes v3.2.7a (25). Posterior probabilities were reported as supporting values for nodes in the trees and scale bar represents substitutions per nucleotide site. Note, that only clade 3 is shown expanded in this figure.

### *In planta* confirmation of diterpene biosynthetic activities of *E. peplus BAHD* genes using Virus Induced Gene Silencing (VIGS)

To further corroborate the function of these identified acyltransferases, VIGS (as previously established for *Euphorbia peplus* (18)) was used to silence expression of the four BAHD genes shown to have activity on ingenol (**5**) or ingenol-3-angelate (**1a**). To avoid off-site targeting, low homology regions of *EpBAHD-06, EpBAHD-07, EpBAHD-08* and *EpBAHD-11* were selected and cloned into a pTRV2 vector containing previously described silencing marker *EpCH42* (18). *Agrobacterium tumefaciens* mediated infiltration was then carried out in three batches: 1), targeting *EpCH42:EpBAHD-06* and *EpCH42:EpBAHD-08;* 2), targeting *EpCH42:EpBAHD-07* and *EpCH42:EpBAHD-11*; and 3), a “double construct” targeting *EpCH42:EpBAHD-07:EpBAHD-11* simultaneously. A construct silencing *EpCH42* independently was used as a control for each experiment.

Chlorotic parts of stems and leaves were harvested around 40 days post-infiltration. Metabolite and transcript levels were then analyzed by LC-MS and qRT-PCR respectively. Transcript levels of both *EpBAHD-08* and *EpBAHD-06* were significantly reduced (3-fold) in stems compared to *EpCH42* – infiltrated controls (Supplementary Figure S11A). However, expression of both BAHD genes was not significantly altered in leaves (Supplementary Figure S11B). Notably, all four *BAHDs* subjected to VIGS were expressed at low levels in leaves (10-20 fold-lower than stem), making detection of subtle changes in gene expression by qRT-PCR unreliable (Supplementary Figure S11B). Next, we analyzed silencing effects on metabolite levels of ingenol-3-angelate (**1a**), ingenol-3-angelate-20-acetate (**2**), and ingenol (**5**), as well as a number of previously isolated ingenane and jatrophane diterpenoids (metabolites **11**-**18**; supplementary Figure S4) (18). Metabolite profiling clearly showed that silencing of *EpBAHD-08* significantly decreased the levels of ingenol-3-angelate (**1a**) and ingenol-3-angelate-20-acetate (**2**) in leaves and stems of *E. peplus*, with the effect being more pronounced in stems where diterpenoid concentrations are higher (Figure 4, Dataset S1). Silencing of *EpBAHD-08* has also led to the accumulation in stem material of ingenol (**5**), a substrate for the *EpBAHD-08* catalyzed reaction (Figure 4, Dataset S1). VIGS thus corroborates the results obtained upon expression of *EpBAHD-08* both *in vitro* and in *N. benthamiana* (Figure 2B and D) and confirms the function of *EpBAHD-08 in planta*, namely the angeloylation of ingenol (**5**) at the C3 position (Figure 1B) to produce ingenol-3-angelate (**1a**). We thus named *EpBAHD-08 ingenol-3-angelate synthase* (*I3AS*). Interestingly, the levels of 20-deoxyingenol-3-angelate (**18**), another ingenane diterpenoid, remained unaltered by silencing of *EpBAHD-08* (Figure 4, Dataset S1), strongly suggesting that this compound is not directly derived from ingenol (**5**) and its biosynthesis does not involve *I3AS*.

**Figure 4.**
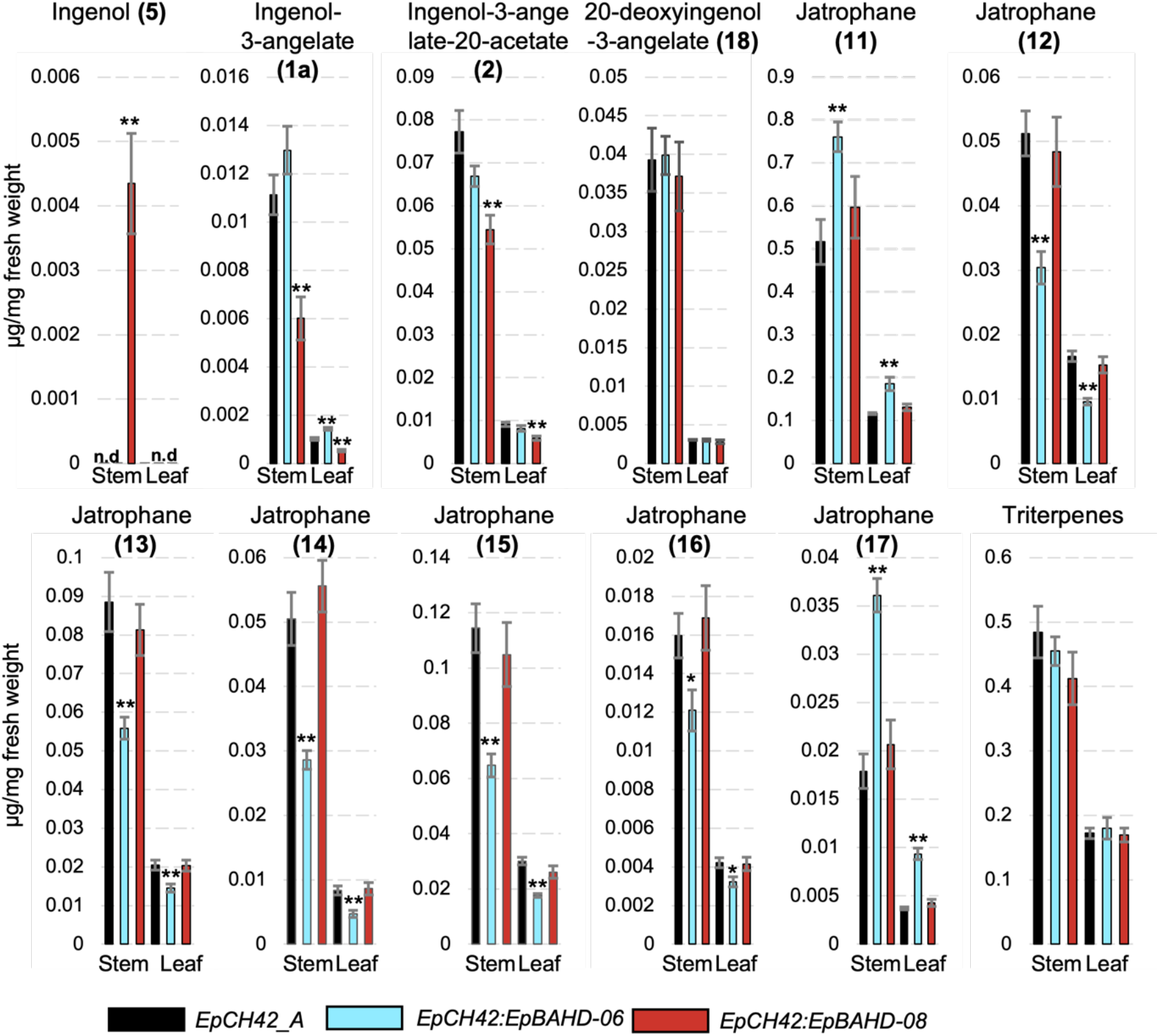
VIGS analysis of *EpBAHD-06 and EpBAHD-08* genes in *Euphorbia peplus*. Metabolite levels in VIGS material were measured for stem and leaves in VIGS marker-only (EpCH42_A, black bars) and marker plus selected BAHD genes: *EpCH42:EpBAHD-06* (cyan bars) and *EpCH42:EpBAHD-08* (red bars). Triterpenes represent the sum of four major triterpenes annotated in Tables S6 and S7. Error bars – SEM (n = 6). Statistically significant (T-test) changes between control (EpCH42_A) and silenced BAHD genes indicated by asterisks separately for each tissue (*-p-value <0.05; ** - p-value <0.01).

In contrast, VIGS of *EpBAHD-06* did not significantly alter the levels of any ingenanes in stems but had a clear effect on the level of jatrophane diterpenoids (Figure 4). *EpBAHD-06*-silenced stems and leaves showed a significant increase of Jatrophane 1 (**11**) and 7 (**17**), accompanied by a strong decrease of Jatrophanes 2-6 (**12-16**) content (Figure 4, Dataset S1). It is noteworthy that the activity of *EpBAHD-06* towards ingenol (**5**) observed in *N. benthamiana* and *in vitro* assays was very low when compared with activity from *EpBAHD-08* (Figure 2B and D), implying it might represent a side-activity for *EpBAHD-06*. The VIGS results showing involvement of *EpBAHD-06* in biosynthesis of jatrophanes rather than ingenanes *in planta* is further confirmation of this.

Silencing of *EpBAHD-07* and *EpBAHD-11, both of* which showed activity on ingenol-3-angelate (**1a**), forming ingenol-3-angelate-20-acetate (**2**) in both *N. benthamiana* and *in vitro* assays, did not alter levels of any of the ingenanes in leaf or stem tissue (Supplementary Figure S12, Dataset S2). qRT-PCR analysis showed only modest (1.5-1.8 fold) transcript reduction in stem tissue (Supplementary Figure S11C and Supplementary Figure S11D). Due to the redundant activity shown for *EpBAHD-07* and *EpBAHD-11*, a separate experiment was performed in an attempt to silence both genes simultaneously. Despite the stronger (2-fold) and consistent reduction in *EpBAHD-07* and *EpBAHD-11* transcript levels in stem tissue (Supplementary Figure S11E and Supplementary S11F), we did not observe any significant effect on level of ingenanes (Supplementary Figure S12, Dataset S3). Levels of some jatrophanes were slightly reduced in *EpBAHD-11 –* silenced stem tissues (Supplementary Figure S12, Dataset S2) but this effect was not seen when *EpBAHD-07* and *EpBAHD-11* were silenced simultaneously (Supplementary Figure S12, Dataset S3), despite EpBAHD-11 transcript levels being more strongly reduced in the latter experiment (Supplementary Figure S11E and Supplementary Figure S11F). Levels of triterpenes also remained unaltered in *EpBAHD-07* and *EpBAHD-11* VIGS experiments (Supplementary Figure S12; Dataset S2 and Dataset S3).

As our VIGS results did not confirm that *Ep*BAHD-07/*Ep*BAHD-011 together are essential for production of ingenol-3-angelate-20-acetate (**2**) *in planta* we assumed there might be further genetic redundancy at this catalytic step in *E. peplus*. Upon closer inspection of our RNAseq data we identified a close homologue of *Ep*BAHD-07 (Supplementary Figure S13; hitherto referred to as *EpBAHD-012*), that could potentially also be involved in the formation of **2**. Indeed, expression of *EpBAHD-12* in *N. benthamiana* also led to acetylation of ingenol-3-angelate (**1a**) at C-20, to produce ingenol-3-angelate-20-acetate (**2**) (Supplementary Figure S14). However, expression levels of *EpBAHD-12* were unaltered in the *EpBAHD-11* – silenced tissues, with its transcript levels being an order of magnitude lower than that of *EpBAHD-07* (Supplementary Figure S11 C-F). Lack of the *in planta* effect on ingenol-3-angelate-20-acetate (**2**) levels seen when *EpBAHD-07* and *EpBAHD-11* were silenced simultaneously could be explained by even greater functional redundancy in the *Euphorbia peplus* genome. In this regard, it is interesting to note that the *BAHD* gene family dramatically expanded during the evolution of land plants, with typically 50-200 BAHD copies present per genome (22). This expansion has resulted in significant functional diversification, as witnessed by the presence of all seven sub-clades of BAHDs in the genomes of angiosperms (22). It is therefore possible that additional functional homologues of *EpBAHD-07, - 11* and *-12* are encoded in the *E. peplus* genome (1). Such a high degree of redundancy would make VIGS challenging. Moreover, 2-3 fold reductions of gene expression might not be sufficient to show a metabolite phenotype.

## Conclusion

Here we report the discovery and characterization of the first genes linked to the committed steps of ingenol-3-angelate (**1a**) biosynthesis in *Euphorbia peplus*. We identified two genes, the products of which catalyze the addition of angelyl-CoA (**9a**) to the ingenol (**5**) scaffold to produce ingenol-3-angelate (**1a**). We demonstrate using VIGS that just one of these genes, EpBAHD-08, is essential for this angeloylation in the native plant *E. peplus*. We also identified three genes whose products can catalyze acetylation of ingenol-3-angelate (**1a**) to ingenol-3-angelate-20-acetate (**2**). In this case, VIGS indicates considerable functional redundancy in the *E. peplus* genome of genes encoding this enzymatic step. Silencing or knockout of these genes would enhance production of ingenol-3-angelate (**1a**) in-planta or give rise to non-acylated compounds, providing enhanced opportunity for semisynthesis of other biologically active ingenol-type diterpenes. Notably, the steps leading to the formation of ingenol (**5**) from the putative intermediate jolkinol C (**7**) remain unclear. The discovery of the late steps of the ingenol biosynthetic pathway now provide a foundation for further discovery efforts in this pharmacologically important class of compounds.

## Materials and Methods

Comprehensive descriptions of materials and methods employed in all experiments are included in the SI Appendix, Material and Methods.

### Plant Material and Growth

*E. peplus* plants were grown in climate chambers (12 h light/12 h dark photoperiod). Plants were kept from 12 pm – 7 pm at 24 °C (50-55 % relative humidity) and during night at 22 °C (60 % relative humidity). Eight-week-old plants were used for transcriptomic and metabolomic studies. *Nicotiana benthamiana* plants were cultivated as recorded previously (23).

### *De Novo* Transcriptome Assembly and Gene Candidate Identification

Total RNA was isolated both from bulk plant tissues using commercially available kits and procedures. Three biological replicates of each tissue were used. Standard mRNA library preparation, Illumina 2×150 bp sequencing and *de novo* transcriptome assembly as well as functional annotation was performed by *BGI Genomics*. Acyltransferase genes were identified by co-expression with the known gene casbene synthase (GenBank: KJ026362.1; Pearson correlation coefficient r ≥ 0.9).

### Agrobacterium tumefaciens-mediated transient transformation of N. benthamiana

Transient expression of acyltransferase gene candidates in tobacco leaves was performed as recently reported (23). Each candidate was tested at least two times in two biological replicates.

### Heterologous expression of candidate genes in *E. coli* and in vitro assays

Genes with acyltransferase activity were recombinantly produced in *E. coli* as previously reported (23). In vitro assays contained 2 µg recombinant protein, 200 µM of the respective CoA donor and 100 µM ingenol, ingenol-3-angelate or ingenol-3-angelate-2-acetate in 25 mM phosphate buffer (pH 7.5). After incubation the assays were stopped by addition of MeOH and filtered solutions were analyzed by liquid chromatography-mass spectrometry.

### Virus-induced gene silencing of acyltransferase genes

VIGS was performed as recently reported (18). The Chlorota 42 marker gene (*EpCH42*) was used as control.

## Data Availability

Gene sequence data has been deposited in GenBank (gene accessions PQ801599-PQ801610). RNAseq data are available as GenBank BioProject PRJNA1214814. All other study data are included in the article and/or supporting information.

## Acknowledgments

We gratefully acknowledge Sarah Heinicke and Maritta Kunert for expert assistance with mass spectrometry. Eva Rothe and the MPI-CE greenhouse team are kindly thanked for the cultivation of *Euphorbia* plants. Dr. Maite Colinas is thanked for helpful discussion. Dr. Moonyoung Kang is thanked for handling of NGS data. CS was generously supported by the Deutsche Forschungsgemeinschaft (DFG; German Research Foundation; Project number: 506268802). Plant art & depiction of microcentrifuge tube in Scheme 2B-F are from biorender.com.

## Author Contributions

CS, PS, IAG and SOC designed the study. CS and MF performed all experiments, except those performed by RMA, TC and AG. RMA synthesized angely- and tiglyl-CoA. TC designed, performed and analyzed the VIGS experiments, helped by AG. YL performed phylogenetic analysis of the BAHD gene family. JW helped with molecular cloning. CS, TC, IG and SOC wrote the manuscript. All authors read and agreed on the final version of this manuscript.

