## Supporting Information for "BAHD news from *Euphorbia peplus*: identification of acyltransferase enzymes involved in ingenane diterpenoid biosynthesis"


**This PDF file includes:**

Supporting text
Figures S1 to S14
Tables S1 to S3
Legends for Datasets S1 to S3
SI References

**Other supporting materials for this manuscript include the following:**

Datasets S1 to S3

### Supporting Information Text

#### Plants and Plant Growth

*Euphorbia peplus* plants were germinated from seeds and grown in climate chambers following a 12 h light/12 h dark photoperiod. Plants were kept from 12-19 h at 24 °C (50-55 % relative humidity) and during night at 22 °C (60 % relative humidity). After eight weeks plants were dissected with a scalpel and tissue samples were snap frozen in liquid nitrogen and stored at – 80 °C until further usage. Identical tissue samples were used both for RNA extraction and metabolomics analysis. Tobacco plants (*Nicotiana benthamiana*) were cultivated as recently described (1). For the purpose of leaf infiltration with *Agrobacterium tumefaciens* GV3101 plants were grown for at least three weeks but no longer than four weeks.

#### Chemicals

Chemicals used in this study were purchased from commercial vendors in molecular biology grade (or higher). Authentic standards of ingenol (**5**) and ingenol-3-angelate (**1a**) were purchased from *TCI* and *Sigma Aldrich*. Ingenol-3-angelate-20-acetate (**2**) was previously isolated from *E. peplus* plants, fully characterized by NMR and reported (2).

#### Molecular Biology Kits & Oligonucleotide Primers

Molecular biology kits were purchased from standard commercial vendors and used according to the manufacturer´s instructions. Total RNA from *E. peplus* was extracted with the RNeasy Mini Kit (*Qiagen*), followed by the removal of unwanted DNA with the TURBO DNA-*free*™ Kit (*Thermo Fischer*). Note, that optionally the RNeasy Mini Kit (*Qiagen*) was used again as a final step to obtain highest purity RNA. Total RNA obtained in this way was directly transcribed into cDNA using Superscript$^{TM}$ IV VILO$^{TM}$ master mix (*Thermo Fischer*). All genes reported in this study were amplified from cDNA with Q5$^{®}$ High-Fidelity 2X Master Mix (*New England Biolabs*), purified by agarose gel electrophoresis (1% agarose, 120 V, 40 min) and recovered thereof using the Zymoclean™ Gel DNA Recovery Kit (*Zymo*). Oligonucleotide Primers (Table S2) were obtained from *Sigma Aldrich*. The In-Fusion kit (*Clontech Takara*) was used for molecular cloning and generated plasmids were isolated from bacterial cultures with the Wizard$^{®}$ *Plus* SV Minipreps DNA Purification System kit (*Promega*). DNA sequencing was used to validate all plasmids generated in this study and performed by *Azenta Life Sciences*.

#### Metabolomics on *E. peplus* Tissue

Dissected plant tissue (50-100 mg) was transferred to a 2 mL microcentrifuge tube equipped with 2x tungsten carbide beads (3 mm diameter; *Qiagen*) and homogenized in a tissue lyser for 2 min at 21 Hz. Afterwards, 3x volumes (150-300 µL) of 80% MeOH (supplemented with 0.1% formic acid and 10 µg/mL olivetolic acid as internal standard) were added and tubes were vortexed vigorously for 2 min. After sonication in a water bath (RT, 15 min) tubes were centrifuged at maximum speed for 20 min. The supernatant was filtered through a syringe-filter (0.2 µm), diluted 1:5 with 80% MeOH (supplemented with 0.1% formic acid and 10 µg/mL olivetolic acid as internal standard) and analyzed by HPLC-MS/MS.

#### RNA Sequencing

Total RNA was prepared as described above and sent to *BGI Genomics* for standard mRNA library preparation and Illumina 2x150 bp sequencing (*ca*. 40 M raw paired-end reads/sample). A *de novo* transcriptome was assembled with Trinity (3) by *BGI Genomics*, using cleaned, trimmed reads as input. Transdecoder (https://github.com/TransDecoder/TransDecoder) was run on the de novo assembled transcriptome to identify all coding regions within the transcripts and functional annotation was performed against several functional databases (NR, NT, GO, KOG, KEGG,

SwissProt, and InterPro). Gene expression was measured in fragments per kilobase of transcript per million mapped reads and calculated with RSEM (4).

#### Identification of Acyltransferase Gene Candidates

The commercially obtained *de novo* transcriptome assembly was used for coexpression analysis. A single transcript for casbene synthase (*Ep*CAS; GeneBank accession number: WCJ37107.1) was identified based on homology to the known sequence of *Ep*CAS. Its expression profile was used to calculate pearson correlation coefficients using Microsoft Excel. Genes annotated as acyltransferaes that showed coexpression (Pearson correlation coefficient r ≥: 0.9) were considered as candidates involved in *Euphorbia diterpenoid* biosynthesis and tested *in planta* for acyltransferase activity. See Supplementary Table S1 for genes that were identified using co-expression analysis.

#### Phylogeny

A BAHD phylogenetic analysis was performed using the 12 identified *E. peplus* acyltransferases identified in this work, as well as 104 non-redundant BAHD sequences from: a) a previous phylogenetic analysis performed by D´Auria *et al*. (5); and b) homologous hits with e-values less than $1x10^{-4}$ from a pBLAST search against the Swissprot database. For gene tree construction all sequences were aligned using MUSCLE v3.8.425. The tree was then constructed using Bayesian analyses (MrBayes v3.2.7a) and tree visualization was performed using Figtree v2.4.4 software. Posterior probabilities were reported as supporting values for nodes in the trees and scale bar represents substitutions per nucleotide site. Two fungal BAHDs were used to form an outgroup to all plant BAHDs. BAHD clades were determined based on Moghe *et al.* 2023 (6).

#### Cloning of gene candidates

Molecular cloning was performed as recently reported in (1). Briefly, genes were amplified from cDNA using primers indicated in Supplementary Table S2 For transient expression in *Nicotiana benthamiana* (*vide infra*) genes were cloned into a modified binary 3Ω1-plasmid (7) by In-fusion cloning (*Clontech Takara*). For recombinant protein production in *Escherichia coli*, full length gene sequences were cloned downstream of a $His_6$-Tag into pOPINF. pOPINF was a gift from Ray Owens (Addgene plasmid #26042).

#### Transformation of *Agrobacterium tumefaciens* GV3101 and transient expression of gene candidates in *Nicotiana benthamiana*

*Agrobacterium tumefaciens* GV3101 was transformed as recently reported (1). Transient gene expression in *N. benthamiana* was performed as described in (8), with minor modifications as reported previously (1). All gene candidates were tested at least 2x times, with biological replicates consisting of at least two leaves from two different tobacco plants.

#### Analysis of plant samples

Snap-frozen tobacco leaf tissue (100 mg) was homogenized on a TissueLyser II (Qiagen) at 22 Hz for 2 min. Homogenized sample was mixed with MeOH + 0.1% formid acid (350 µL). After vigorous vortexing (1-2 min) and sonication (RT, 15 min) samples were centrifuged a max. speed (> 13000 x g; 20-30 min). All samples were filtered through 0.22 µM PTFE syringe filters and analyzed by high-resolution LC-MS. For data analysis the software DataAnalysis Version 5.3 (Bruker) was used.

#### Recombinant protein production in *Escherichia coli*

All BAHD-acyltransferase genes were heterologously expressed in chemically competent *E. coli* BL21(DE3). Transformed cells were streaked for single colonies and inoculated into 10 mL of LB medium supplemented with 100 µg/mL carbenicillin. This seed culture was cultivated for 16-20 h at 37 °C and 200 rpm. An aliquot of the seed culture (1 mL/8 mL) was inoculated into 2XYT Medium (100 mL/1000 mL) supplemented with 100 µg/mL carbenicillin or 50 µg/mL kanamycin. The resulting production culture was incubated at 37 °C and 200 rpm until the optical density ($OD_{600}$) reached 0.4-0.6. Acyltransferase expression was induced by adding isopropyl-*β*-D-1-thiogalactopyranoside (IPTG) to a final concentration of 200 µM and cells were cultivated at 18 °C and 200 rpm for up to 24 h. Afterwards the culture medium was removed by centrifugation (4000 rpm; 4 °C; 20-30 min) and the cell pellets were snap-frozen in liquid nitrogen.

**Purification of recombinant proteins**

Cell pellets were thawed on ice and resuspended in 10 mL (small scale purification) or 80-100 mL (large scale purification) of buffer A (50 mM Tris base, 50 mM glycine, 500 mM NaCl, 20 mM imidazole, 5% glycerol (v/v), pH 8.0; a fresh 100 ml aliquot of this was prepared on the day of protein purification and mixed with 10 mg lysozyme and 1x protease inhibitor cocktail tablet [cOmplete, EDTA-free, *Roche*]). Resuspended cells were lysed using an ultrasonic liquid processor (vibra cell™, *Sonics*®; 40 % amplitude; 2 s on/3 s off; total `on´-time: 3 min). Cell debris was removed by centrifugation (4 °C, 35 min, 35000 x g). The acyltransferase proteins were then purified using NiNTA beads (*Takara* Hi60 Superflow Resin) according to manufacturer's instructions for small scale purification or on an ÄKTA pure FPLC system (*GE Healthcare*) connected to a HisTrap™ column (*cytiva*, column volume = 5 mL) for large scale purification. The FPLC system was programed to: [A], equilibrate the column (flow rate = 5 ml/min) with 5x column volumes of buffer A (50 mM Tris base, 50 mM glycine, 500 mM NaCl, 20 mM imidazole, 5% glycerol [v/v], pH 8.0); [B] load the protein sample (flow rate = 2 ml/min); [C] wash the column [flow rate = 5 ml/min] with buffer A until the UV absorption at 280 nm is stable (stability time = 1 min; accepted UV fluctuation = 0.1 mAU; maximum wash volume = 20x column volumes); [D] elute the protein with 5x column volumes of buffer B (50 mM Tris base, 50 mM glycine, 500 mM NaCl, 500 mM imidazole, 5% glycerol (v/v), pH 8.0). Elution of the protein of interest was monitored using UV absorption at 280 nm. Fractions of interest were assessed by SDS gel electrophoresis, pooled and rebuffered to buffer C (20 mM HEPES, 150 mM NaCl, pH 7.5). For small scale purification NiNTA beads were repeatedly washed with buffer A and then eluted with buffer B. Ultimately, buffer B was exchanged for buffer C using Amicon 10 kDa concentrator columns (Merk Millipore).

**Enzymatic *in vitro* assays**

Recombinant BAHD acyltransferases *Ep*BAHD-06 and *Ep*BAHD-08 were tested for acylation activity on ingenol (**5**), assessing both angelyl-CoA (**9a**) and tiglyl-CoA (**9b**) as donors. Reaction mixtures (100 µL total volume, 25 mM $K_2HPO_4/KH_2PO_4$ pH = 7.5) comprised 200 µM CoA donor, 100 µM ingenol (**5**) and 2 µg recombinant protein. Reactions were started by the addition of the CoA donor and then incubated 1 h at 30 °C / 300 rpm, protected from light. Negative controls consisted of proteins purified from cultures expressing the EV. Reactions were quenched by addition of an isovolume of MeOH 100%. Prior to LC-MS analysis the reactions were filtered through a PTFE filter (0.22 µm).

Recombinant BAHD acyltransferases *Ep*BAHD-07 and *Ep*BAHD-11 were tested for acetylation activity on ingenol-3-angelate (**1a**). Reaction mixtures (100 µL total volume, 25 mM $K_2HPO_4/KH_2PO_4$ pH = 7.5) comprised 200 µM acetyl-CoA donor, 100 µM ingenol-3-angelate **(1a)** and 2 µg recombinant protein. Reactions were started by the addition of the protein and then incubated 1 h at 30°C / 300 rpm, protected from light. Negative controls consisted of proteins purified from cultures expressing the EV. Reactions were quenched by addition of an isovolume of MeOH 100%. Prior to LC-MS analysis the reactions were filtered through a PTFE filter (0.22 µm).

**LCMS data acquisition**

All metabolites reported in this study were analysed using method 1 and 2, except for those metabolite studies conducted during VIGS experiments. For VIGS metabolite analysis a different methodology was deployed and is outlined *vide infra.* Instrumentation for method 1 and 2 has been previously reported by us, and only gradient settings were changed (1, 9, 10). In brief, for LCMS data acquisition an impact II UHR-Q-ToF (Ultra-High Resolution Quadrupole-Time-of-Flight) mass spectrometer (Bruker Daltonik; Bremen, Germany) connected to an Ultra-high performance liquid chromatography system Ultimate 3000 RS Thermo Fisher Scientific, Germering, Germany was used. Compound separation was achieved using reverse-phase liquid chromatography on a Phenomenex Kinetex XB-C18 (100 x 2.1 mm, 2.6 µm; 100 Å) column operated at 40 °C. Mobile phases: (A) water with 0.1% formic acid; (B) acetonitrile; flow rate = 0.6 mL/min. 2 µL sample was injected in each run; authentic standards were prepared as methanol solutions in concentration ranges between 20-100 µM. Chromatography conditions method 1: 10% B for 2 min; 10% B to 90% B in 10 min; 90% B to 100% B in 0.2 min for 3.0 min and 10% B for 4.7 min. Chromatography conditions method 2: 10% B for 1 min; 10% B to 90% B in 10 min; 90% B to 100% B in 0.2 min for 3.0 min and 10% B for 2.7 min. Mass spectrometry conditions: mass spectrometry was performed in positive electrospray ionization mode (capillary voltage = 3500 V; end plate offset = 500 V; nebulizer pressure = 2.5 bar; drying gas: nitrogen at 250 °C and 11 L/min). Mass spectrometry data was recorded at 12 Hz ranging from 80 to 1000 m/z using data dependent MS2 and an active exclusion window of 0.2 min. Tandem mass spectrometry settings: fragmentation was triggered on an absolute threshold of 400 and restricted to a total cycle time range of 0.5 s; collision energy was deployed in a stepping option model (20-50 eV) At the beginning of each run, a sodium formate-isopropanol solution was injected by a syringe pump at 0.18 mL $h^{-1}$, and the m/z values were recalibrated using the expected cluster ions. The initial 1 min of the chromatographic gradient was directed towards the waste. Chromatographic method 2 was utilized in Fig. 1C, S9 and S15. All the other figures were generated with method 1.

**NMR Analysis**

Nuclear magnetic resonance (NMR) spectra were recorded on a 400 MHz Bruker Advance III HD spectrometer (Bruker Biospin GmbH, Rheinstetten, Germany) at *ca*. 293 K. Samples for NMR analysis were prepared by dissolving approximately 5 mg of sample in 0.55 mL $D_2O$. Chemical shift values ($\delta_H$) are reported in parts per million (ppm) relative to residual solvent ($\delta_H$ 4.79, s), and coupling constants (*J*) are expressed in hertz (Hz), in the following format: chemical shift value (multiplicity, coupling constant, integration). $^1$H NMR spectral data are described, using the following abbreviations: appbrs (apparent broad singlet), d (doublet), dd (doublet of doublets), m (multiplet), s (singlet), and t (triplet).

**General Procedure: Synthesis of CoA thioesters 9a and 9b.**

**NMR Analysis**

Nuclear magnetic resonance (NMR) spectra were recorded on a 400 MHz Bruker Advance III HD spectrometer (Bruker Biospin GmbH, Rheinstetten, Germany) at *ca*. 293 K. Samples for NMR analysis were prepared by dissolving approximately 5 mg of sample in 0.55 mL $D_2O$. Chemical shift values ($\delta_H$) are reported in parts per million (ppm) relative to residual solvent ($\delta_H$ 4.79, s), and coupling constants (*J*) are expressed in hertz (Hz), in the following format: chemical shift value (multiplicity, coupling constant, integration). $^1$H NMR spectral data are described, using the following abbreviations: appbrs (apparent broad singlet), d (doublet), dd (doublet of doublets), m (multiplet), s (singlet), and t (triplet).

**General Procedure: Synthesis of CoA thioesters 9a and 9b.**

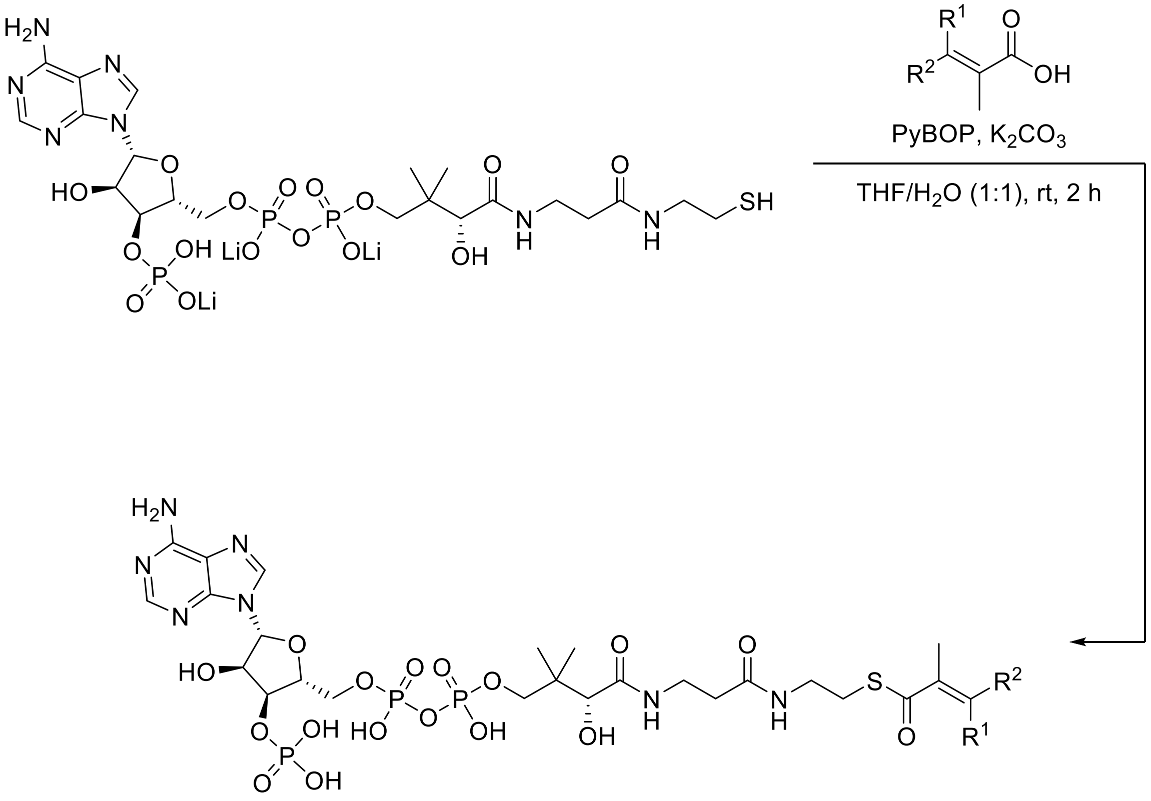

**9a** ($R^1$ = $CH_3$, $R^2$ = H), 26%
**9b** ($R^1$ = H, $R^2$ = $CH_3$), 66%

To an 8 mL vial was added the appropriate carboxylic acid (3.2 mg, 32 µmol), PyBOP (33.3 mg, 64 µmol), and THF/$H_2O$ (1:1, 2.46 mL). The mixture was treated with $K_2CO_3$ (17.55 mg, 127 µmol) and stirred at room temperature for 5 min. A solution of CoA trilithium salt in THF/$H_2O$ (1:1, 0.54 mL) was then added to the reaction mass. After stirring at room temperature for a further 2 h (complete consumption of starting material was confirmed by RP-TLC [20% MeOH in $H_2O$ with 0.1% (v/v) $HCO_2H$]), the resulting mixture was then partially concentrated under reduced pressure and subjected to RP-PTLC (20% MeOH in $H_2O$ with 0.1% [v/v] $HCO_2H$), to afford the corresponding CoA thioesters **9a** and **9b**, respectively, as colorless solids.

Note: Following isolation, CoA thioesters 9a and 9b were stored as solids at – 80 °C.

**Tiglyl-CoA (9b)**

Following the General Procedure for the synthesis of CoA thioesters, using tiglic acid, gave tiglyl CoA as a colorless solid (18 mg, 66%): $^1$H NMR (400 MHz, $D_2O$) $\delta$ 8.53 (s, 1H), 8.23 (s, 1H), 6.83 (qq, $J$ = 6.8, 1.3 Hz, 1H), 6.16 (d, $J$ = 6.1 Hz, 1H), 4.87–4.82 (m, 3H), 4.60–4.59 (m, 1H), 4.27–4.25 (m, 2H), 4.03 (s, 1H), 3.85 (dd, $J$ = 9.7, 5.0 Hz, 1H), 3.58 (dd, $J$ = 9.8, 4.9 Hz, 1H), 3.45 (t, $J$ = 6.5 Hz, 2H), 3.33 (t, $J$ = 6.9 Hz, 2H), 2.98 (t, $J$ = 6.4 Hz, 2H), 2.43 (t, $J$ = 6.5 Hz, 2H), 1.80–1.77 (m,

6H), 0.91 (s, 3H), 0.77 (s, 3H); HRMS (ESI/Q-TOF) m/z: [M−H]⁻ Calcd. for $C_{26}H_{41}N_7O_{17}P_3S$ 848.1498; Found 848.1502. Spectral data was consistent with that previously reported.(11)

$^{1}H$ NMR ($D_2O$, 400 MHz) Tiglyl-CoA (9b)

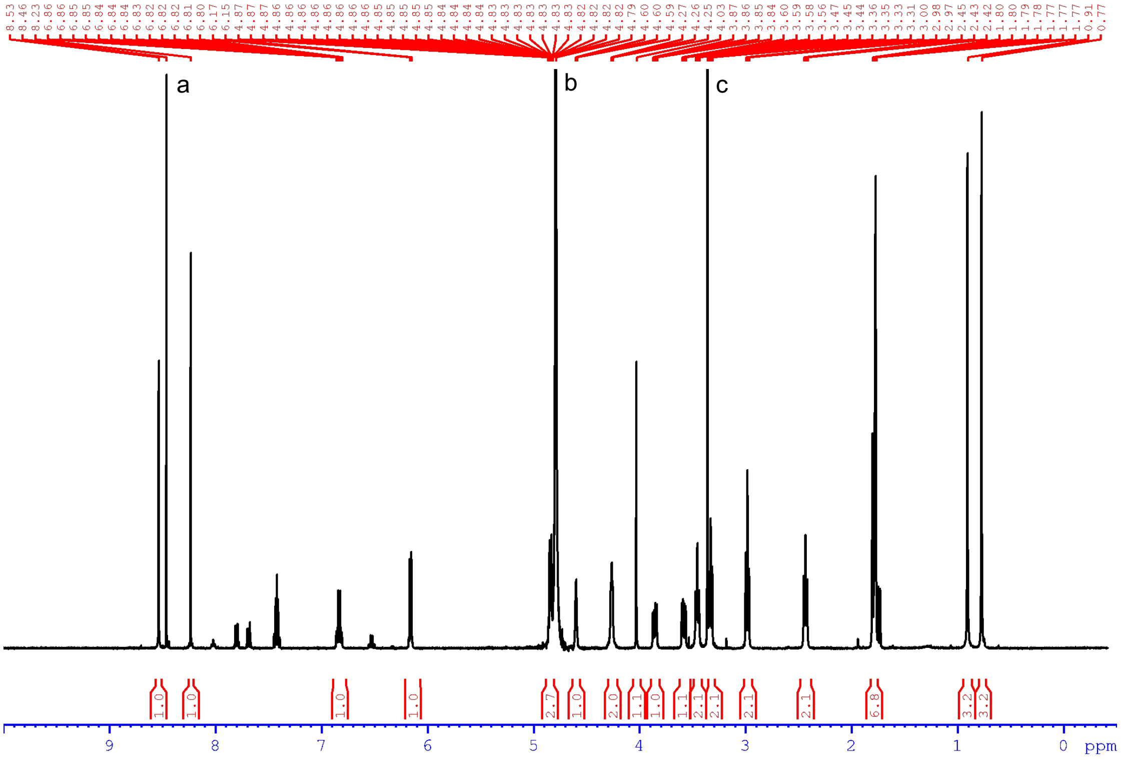


[a]$HCO_2H$;[b]$H_2O$;[c]MeOH.

**Angelyl-CoA (9a)**

Following the General Procedure for the synthesis of CoA thioesters, using angelic acid, gave angelyl CoA as a colorless solid (7 mg, 26%):$^{1}H$ NMR (400 MHz, $D_2O$) $\delta$ 8.54 (s, 1H), 8.25 (s, 1H), 6.87–6.82 (m, 1H), 6.13 (d, $J$ = 6.0 Hz, 1H), 4.85–4.82 (m, 3H), 4.60 (appbrs, 1H), 4.26 (appbrs, 1H), 4.03 (s, 1H), 3.86 (dd, $J$ = 9.8, 4.5 Hz, 1H), 3.58 (dd, $J$ = 9.7, 4.8 Hz, 1H), 3.46 (t, $J$ = 6.4 Hz, 2H), 3.33 (t, $J$ = 6.5 Hz, 2H), 2.99 (t, $J$ = 6.4 Hz, 2H), 2.44 (t, $J$ = 6.4 Hz, 2H), 1.81–1.78 (m, 6H), 0.91 (s, 3H), 0.78 (s, 3H); HRMS (ESI/Q-TOF) m/z: [M−H]⁻ Calcd. for $C_{26}H_{41}N_7O_{17}P_3S$ 848.1498; Found 848.1499.

**$^1$H NMR ($D_2O$, 400 MHz) Angelyl-CoA (9a)**

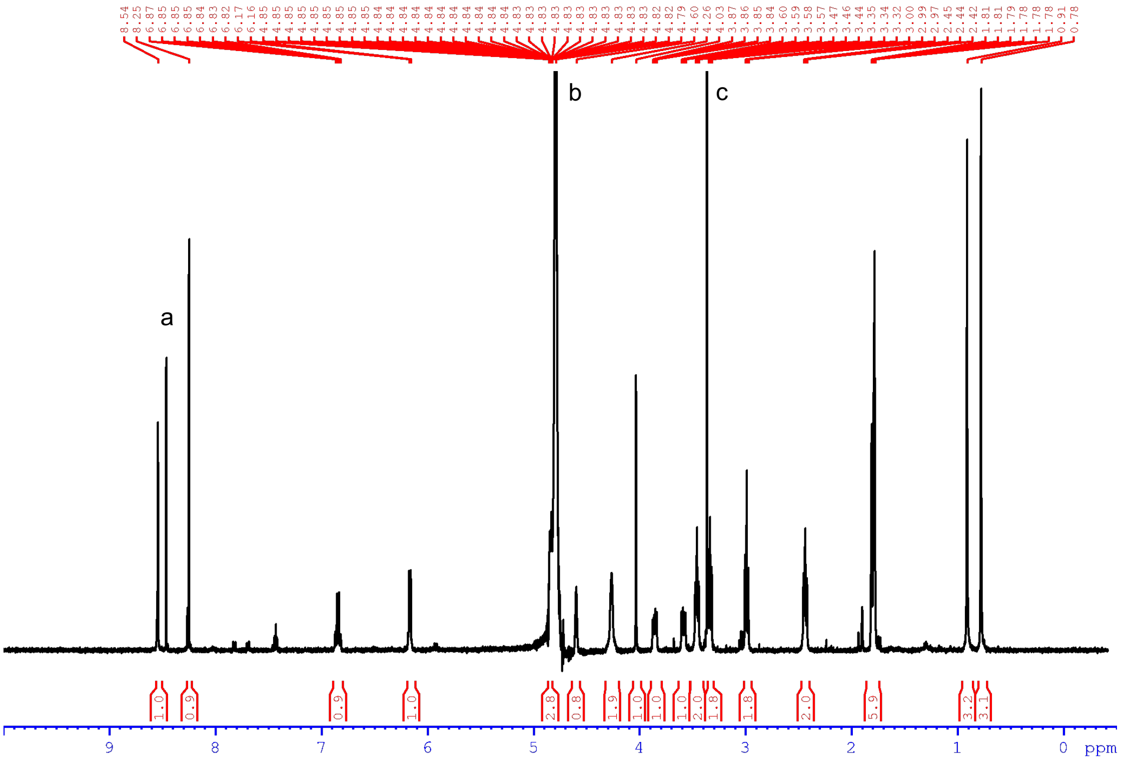


[a]$HCO_2H$;[b]$H_2O$;[c]MeOH.

**Virus-induced gene silencing (VIGS) of: *EpCH42, EpBAHD-06, EpBAHD-07, EpBAHD-08 and EpBAHD-11.***

Experimental design followed previously published (2) VIGS experiments established for *Euphorbia peplus* using Chlorota 42 marker gene (*EpCH42*). A 165bp, 173bp, 174bp and 124bp fragments from the non-conserved regions (*C*-terminus) of the *EpBAHD-06, EpBAHD-07, EpBAHD-08* and *EpBAHD-11,* respectively, were PCR-amplified from *E. peplus* stem cDNA using *XhoI* and *SmaI* In-Fusion-tailed PCR primers (Table S2). The amplified fragments were inserted into the pTRV2-*EpCH42*-Vigs construct (2) digested with *XhoI* and *SmaI* via In-Fusion cloning tools (TaKaRa bio Inc. Kusatsu, Japan), according to the manufacturer's protocol to form the pTRV2-*EpBAHD-06:EpCH42*-Vigs, pTRV2-*EpBAHD-07:EpCH42*-Vigs, pTRV2-*EpBAHD-08:EpCH42*-Vigs and pTRV2-*EpBAHD-11:EpCH42*-Vigs constructs. Construct pTRV2-*EpBAHD-07:EpBAHD-11:EpCH42*-Vigs for simultaneous silencing of *EpBAHD-08* and *EpBAHD-11* was created by amplifying synthetic fragment containing fusion of 174 and 124bp fragments described above with *XhoI* and *SmaI* In-Fusion-tailed PCR primers (Table S2) and inserting it into the pTRV2-*EpCH42*-Vigs construct (2) as described above.

After confirming the presence of the correct inserts by Sanger sequencing, the pTRV2 vectors were transformed into *A. tumefaciens* GV3101 using the freeze-thaw method (21). The *A. tumefaciens* GV3101 strains containing pTRV1, pTRV2-*EpCH42*-Vigs and one of the five target-gene constructs (*EpBAHD-06, EpBAHD-07, EpBAHD-08, EpBAHD-11* and *EpBAHD-11:EpBAHD-07*) were grown separately overnight at 28 °C, 220 rpm, in 10 mL LB medium containing kanamycin and gentamycin (50 mg/L) antibiotics. A 1 ml aliquot of overnight-grown cultures was inoculated in 50 mL of LB medium containing 10 mM MES and 20 µM acetosyringone with kanamycin and gentamycin (50 mg/L) antibiotics and grown overnight at 28 °C, 220rpm. *A. tumefaciens* cells were harvested and re-suspended in the infiltration buffer (10 mM $MgCl_2$, 10 mM MES, pH 5.6, 200 µM acetosyringone) to a final $OD_{600}$ of 2.5 (for both pTRV1 and pTRV2 and its derivatives) and shaken for 2 h at 28 °C, 100 rpm. Equal volumes of the *A. tumefaciens* cultures carrying one of the pTRV2-derived constructs were mixed with pTRV1-carrying cultures. Mixed *A. tumefaciens* cultures were infiltrated into both cotyledons of *E. peplus* seedlings 9-days after sowing, using a 1 mL syringe. Infiltrated plants were grown under 16 h / 8 h light and 25 °C / 22 °C day/night regime. Around 150-160 plants were infiltrated for pTRV2-*EpCH42*-Vigs control group and for each of pTRV2-*EpCH42*-Vigs derived target gene constructs.

Chlorotic parts of leaf and stem samples were collected separately from plants 6 weeks post-infiltration. Fresh plant material was pooled from 6-8 independent plants to form one biological replicate, flash frozen in liquid $N_2$ and stored in - 80 °C. Five or six biological replicates were used for each of the two groups: pTRV2-*EpCH42*-Vigs controls and one of the three pTRV2-*EpCH42*-Vigs derived target gene constructs. Experiment was done in three batches, first one contained pTRV2-*EpBAHD-06:EpCH42*-Vigs, pTRV2-*EpBAHD-08:EpCH42*-Vigs and pTRV2-*EpCH42*-Vigs control, second one contained: pTRV2-*EpBAHD-07:EpCH42*-Vigs, pTRV2-*EpBAHD-11:EpCH42*-Vigs and pTRV2-*EpCH42*-Vigs control and third one contained pTRV2-*EpBAHD-07:EpBAHD-11:EpCH42*-Vigs and pTRV2-*EpCH42*-Vigs control constructs.

**Metabolite and mRNA transcript analysis of VIGS treated *E. peplus* plants.**

Plant material was ground in liquid nitrogen using a mortar and pestle. 300-330 mg of the ground fresh tissue was extracted using ethyl acetate with PMA internal standard and run on LC-MS as described before (Czechowski et al. 2022).

50-100 mg of the same ground fresh tissue was used to extract total RNA from stem tissue using the RNAeasy kit (Qiagen, Hilden, Germany) with "on column" DNAse digestion according to manufacturer's instructions. RNA was precipitated overnight using 0.1 volume of 3 M sodium acetate and 2.5 volumes of 100% ethanol, washed twice the following day with 70% ethanol, air dried and re-suspended in water. Total RNA was extracted from leaf tissue using the CTAB-lithium chloride method (20) RNA samples were DNase treated and further purified using the on-column digestion protocol for the Qiagen RNeasy miniprep kit. cDNA was synthesized from 5 µg of total RNA using random hexamers using Superscript II reverse transcriptase (Life Technologies). qPCR primers (Table S2) were designed using Geneious Prime® 2022.2.1 software. Real-time PCR was performed on CFX384 Real-Rime System (Bio-Rad Laboratories) using iTAQ™ SYBR® Green Supermix (Bio-Rad Laboratories). Each 10 µL reaction contained 2 µL of a 3-fold dilution of the cDNA synthesis reaction, 5 µL of 2X Supermix, and primers at a final concentration of 250 nM. The cycling conditions included an initial activation step for 5 s at 95 °C followed by 40 cycles of denaturation at 95 °C for 10 s and annealing/extension at 60 °C for 30 s. Fluorescence data were acquired during the annealing/extension phase. A melt curve was obtained at the end of the amplification to allow confirmation of product specificity. $C_T$ values were obtained using CFX Manager Software (Bio-Rad laboratories) and amplification efficiencies (E) obtained using LinReg PCR (21). Transcript abundance for the gene of interest (GOI) relative to housekeeping gene (HKG) was determined using the formula: GOI expression level = $2^{\Delta C_T} / 2^{\Delta C_T}$. Housekeeping genes were selected based on previously used for gene quantification by qRT-PCR in *Euphorbia peplus* (Czechowski et al. 2022): a homologue of *Elongation Factor 1α* (*EpEF1α;* gene_bank accession OL744076) and a homologue of *SUMO-conjugating enzyme SCE1* (*EpSumo,* gene_bank accession OL744078).

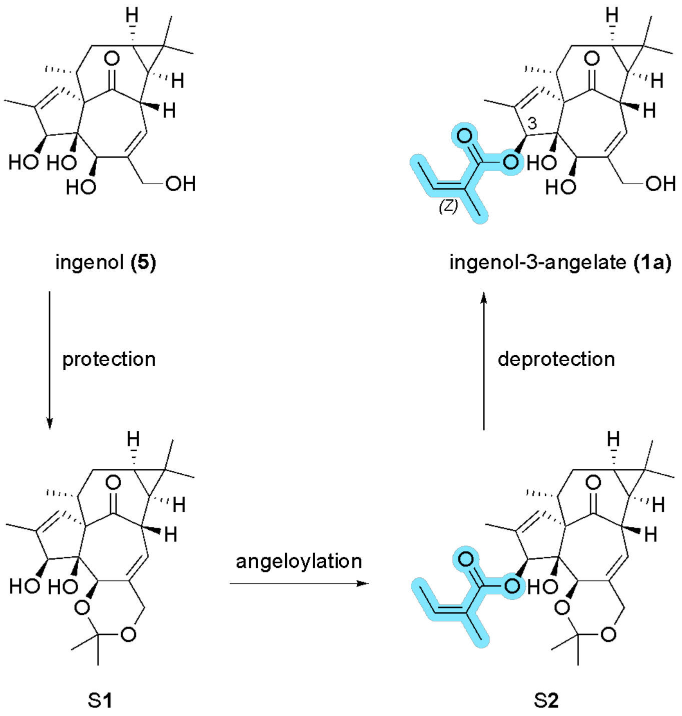


**Fig. S1.** Chemical synthesis of ingenol-3-angelate (**1a**) from ingenol (**5**) as reported by Liang *et al.* in 2012 (12). While **5** is more abundantly produced *in planta* than **1a** it has to be extracted from seeds of *Euphorbia lathyris* (250 mg per kg of dried seeds) (13).

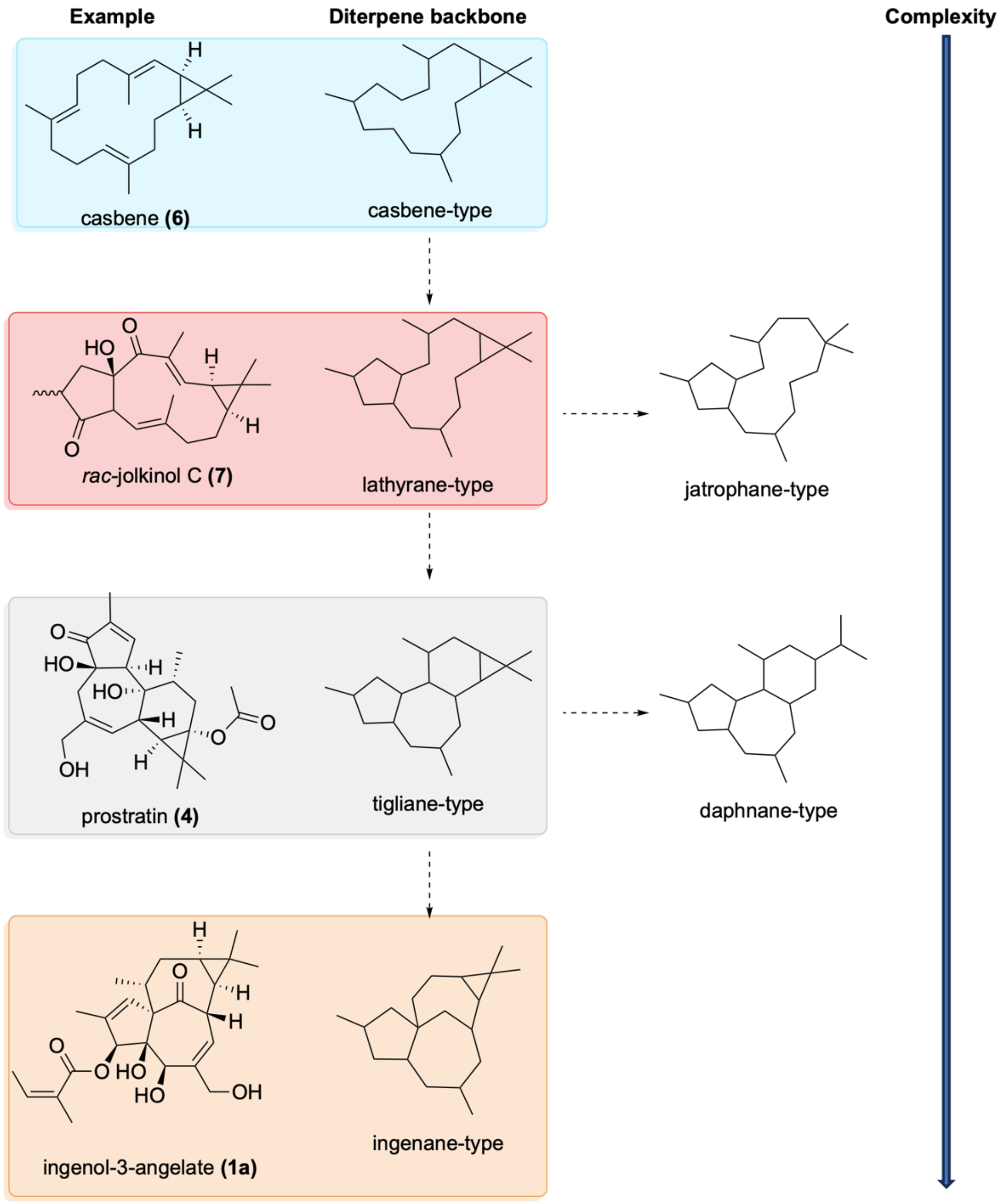


**Fig. S2.** Structures of the major *Euphorbia* diterpene scaffolds jatrophane, lathyrane, tigliane, daphnane and ingenane.

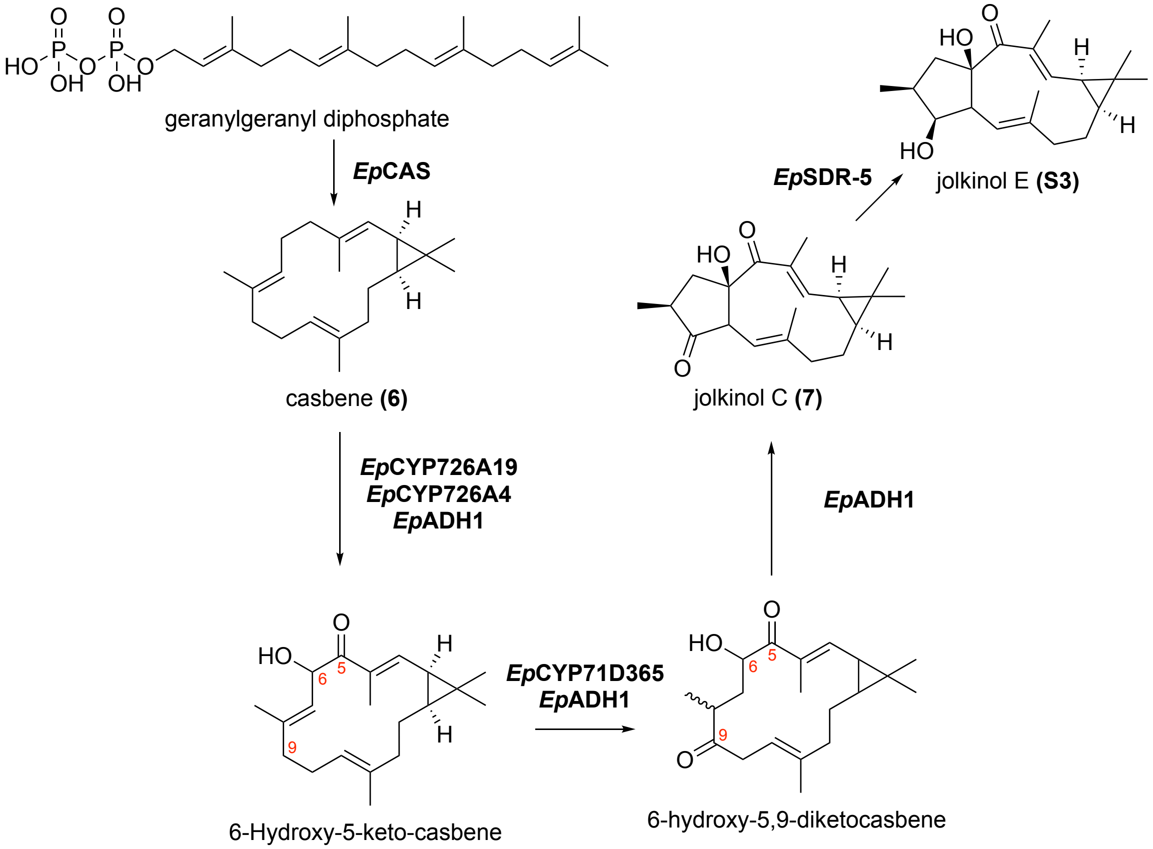


**Fig. S3.** Biosynthesis of jolkinol C (**7**) and jolkinol E (**S3**) in *E. peplus*. Genes first reported by Luo *et al.* 2016 and Czechowski *et al.* 2022 (2, 14).

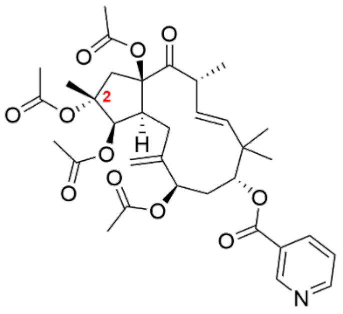

Jatrophane 1 **(11)**

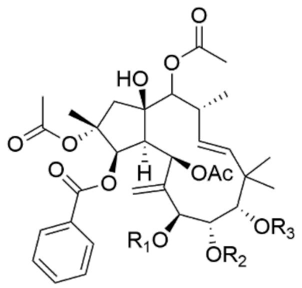

Jatrophane 2 **(12)** $R_1$= Ac; $R_2$ = Ac; $R_3$ = Ac
Jatrophane 3 **(13)** $R_1$= iBu; $R_2$ = H; $R_3$ = Nic
Jatrophane 4 **(14)** $R_1$= Ac; $R_2$ = H; $R_3$ = Nic
Jatrophane 5 **(15)** $R_1$= Ac; $R_2$ = Ac; $R_3$ = Nic
Jatrophane 6 **(16)** $R_1$= Ac; $R_2$ = H; $R_3$ = Ac

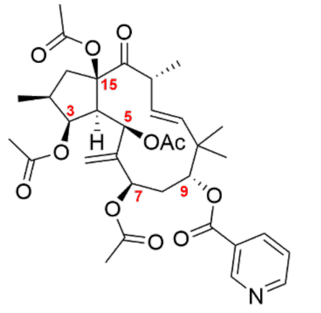

Jatrophane 7 **(17)**

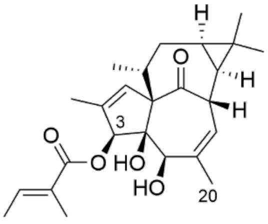

20-deoxingenol-3-angelate (**18**)

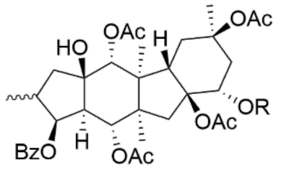

**S4** R = H
**S5** R = Ac

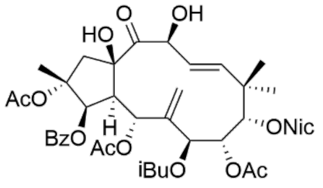

**S6**

**Fig. S4.** Examples of (multi)acylated diterpenoids from *Euphorbia peplus* (15–18).

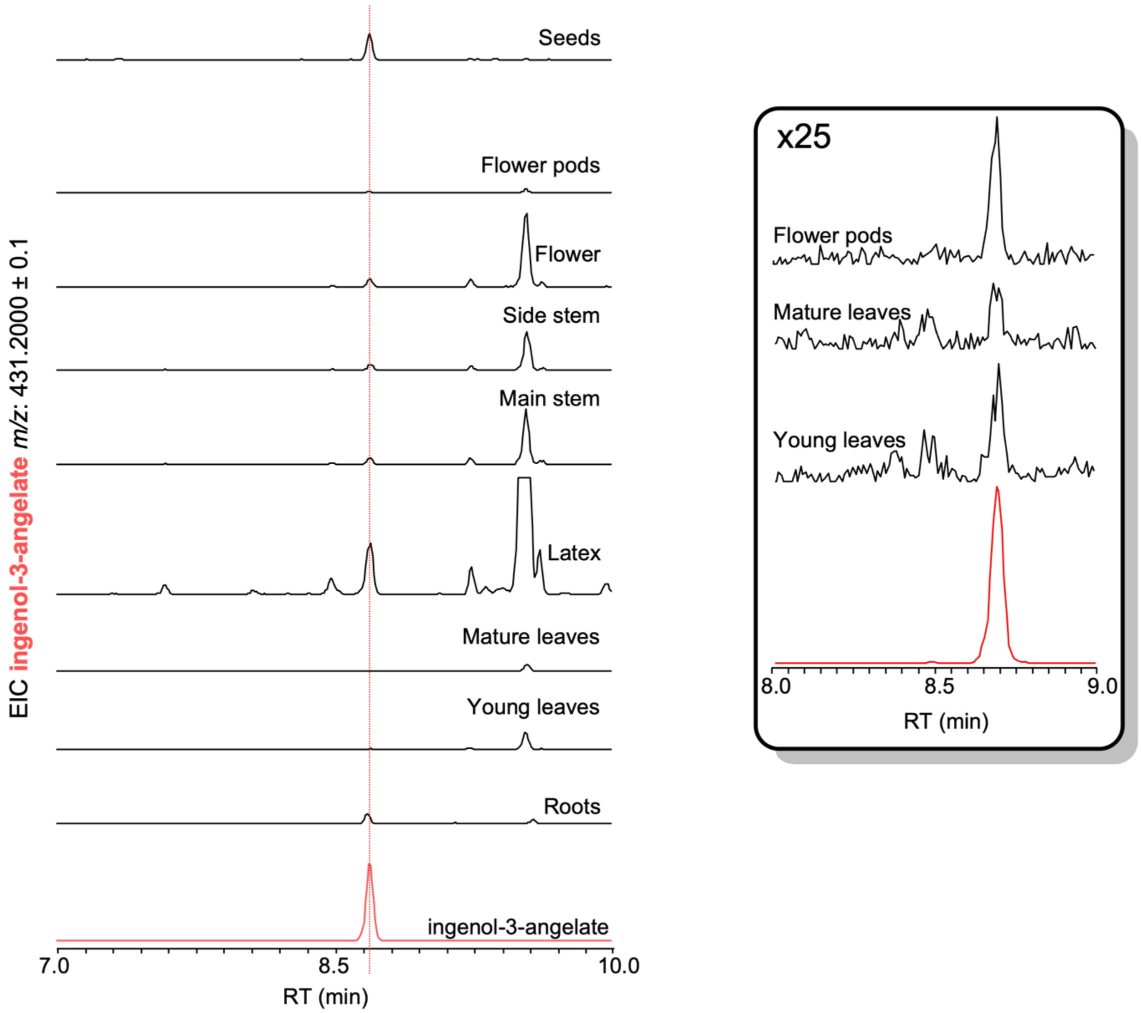


**Fig. S5.** Observed levels of ingenol-3-angelate (**1a**) in different tissues of *E. peplus*. Methanolic extracts were analyzed using HPLC-MS method 1. Extracted ion chromatograms (EIC; $[M+H]^+$ = 431.2) were compared to an authentic standard of **1a** (bottom trace, red). The identification of **1a** was confirmed through matching retention times and MS/MS data. For flower pods, mature leaves, and young leaves, the extracted ion chromatograms of ingenol-3-angelate (**1a**) were magnified 25x (see box on the right) to enhance visibility.

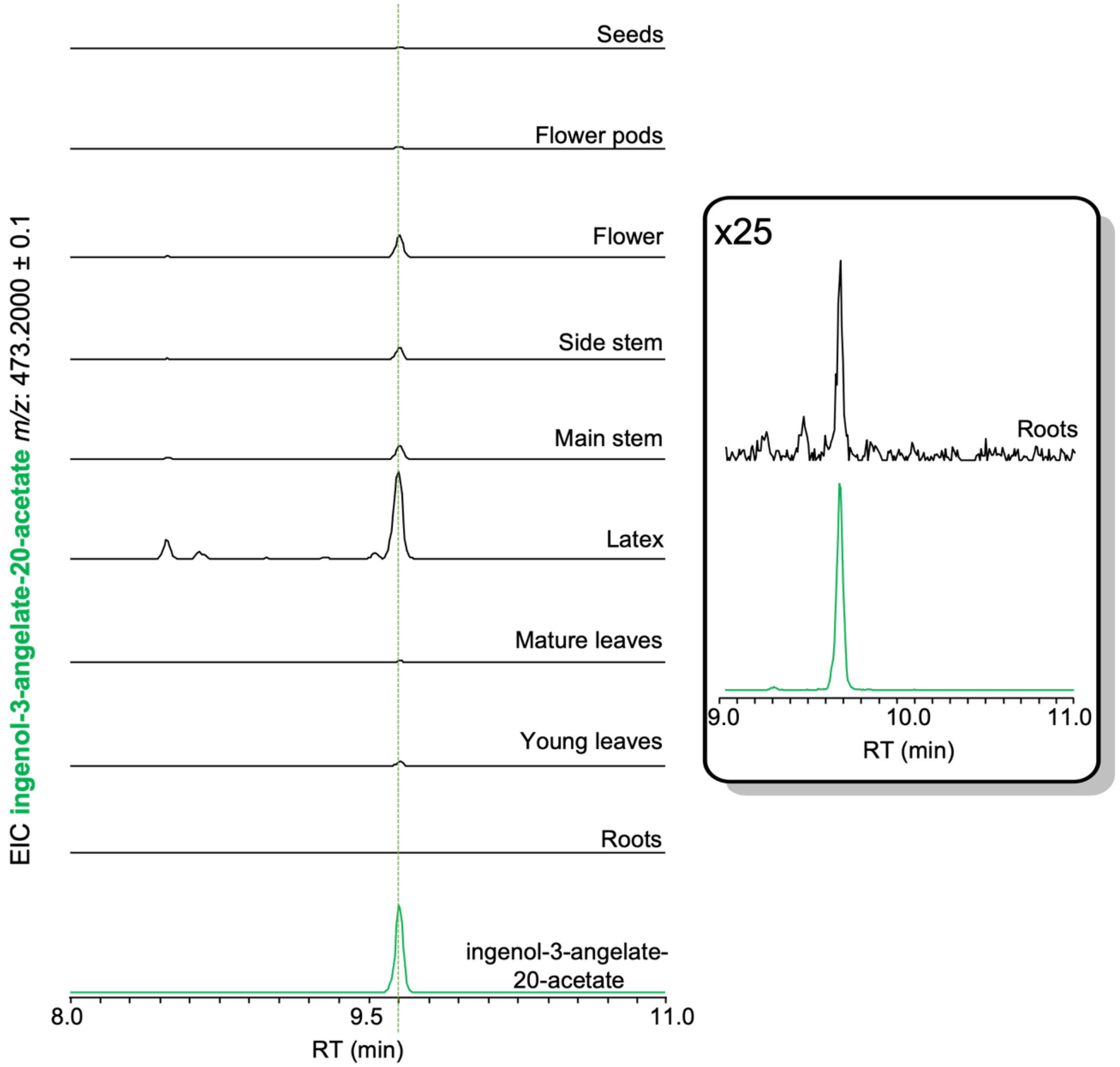


**Fig. S6.** Observed levels of ingenol-3-angelate-20-acetate (**2**) in different tissues of *E. peplus*. Methanolic extracts were analyzed using HPLC-MS method 1. Extracted ion chromatograms (EIC; $[M+H]^+$ = 473.2) were compared to an authentic standard of **2** (bottom trace, green). The identification of **2** was confirmed through matching retention times and MS/MS data. For root tissue the extracted ion chromatograms of ingenol-3-angelate-20-acetate (**2**) were magnified 25x (see box on the right) to enhance visibility.

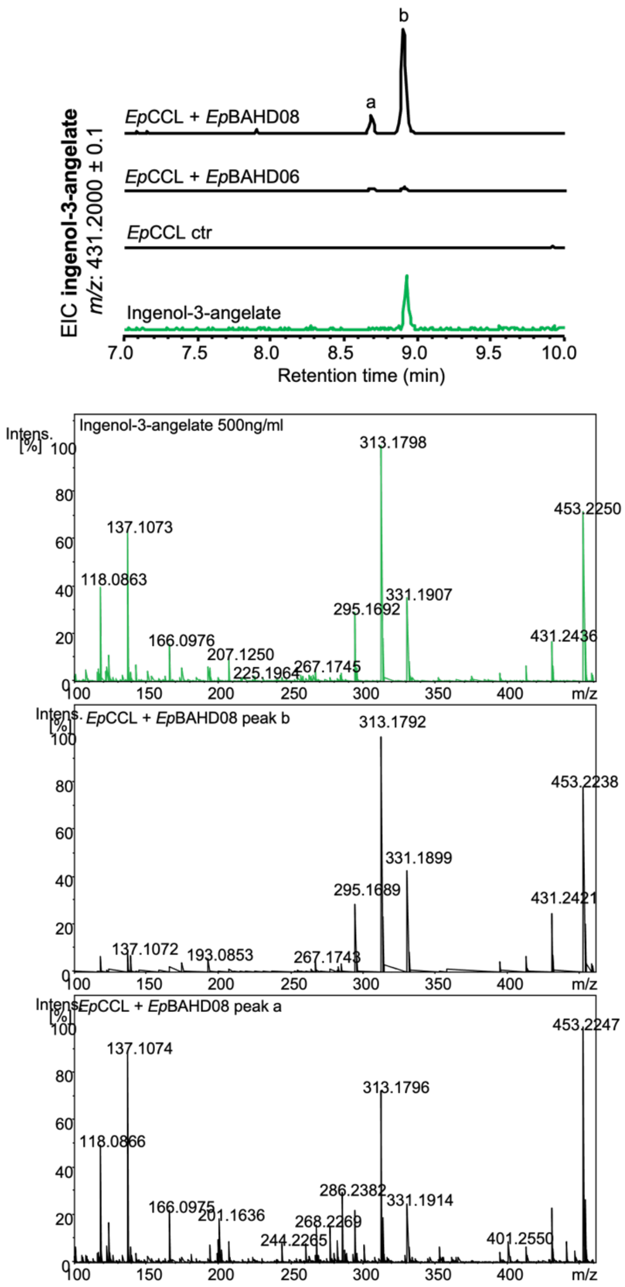


**Fig. S7.** Fragmentation pattern analysis (MSMS) of ingenol-3-angelate (**1a**; peak b) and unknown product **1b** (peak a) using an authentic standard of **1a** for comparison. MSMS patterns are identical, leading to the tentative identification of **1b** as ingenol-3-tigliate.

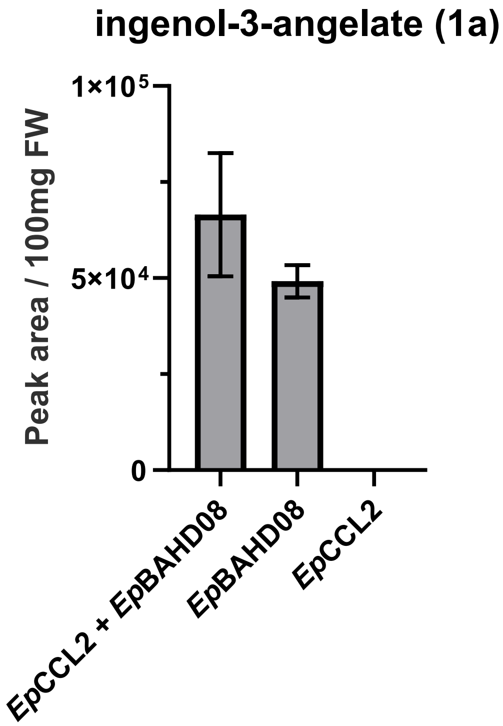


**Fig. S8.** Angelyl-CoA-ligase (*Ep*CCl2) is not required for product formation in *N. benthamiana*. Infiltration of *Ep*BAHD-08 together with *Ep*CCL2 did not lead to significantly improved levels of ingenol-3-angelate (**1a**).

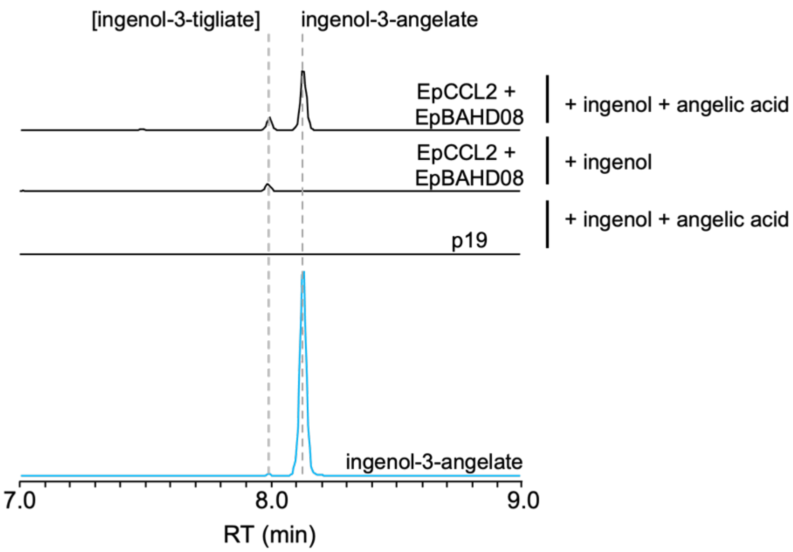


**Fig. S9.** Angelic acid is required for product formation in *N. benthamiana*. Formation of ingenol-3-angelate (**1a**) was only observed when *N.* benthamiana leaves were co-infiltrated with ingenol, angelic acid as well as *EpCCL2* and *EpBAHD-08* (top trace). In the absence of angelic acid (expression of *EpCCL2* and *EpBAHD-08* in the presence of ingenol) no formation of ingenol-3-angelate (**1a**) was observed. Infiltration of p19 on its own was used as negative control.

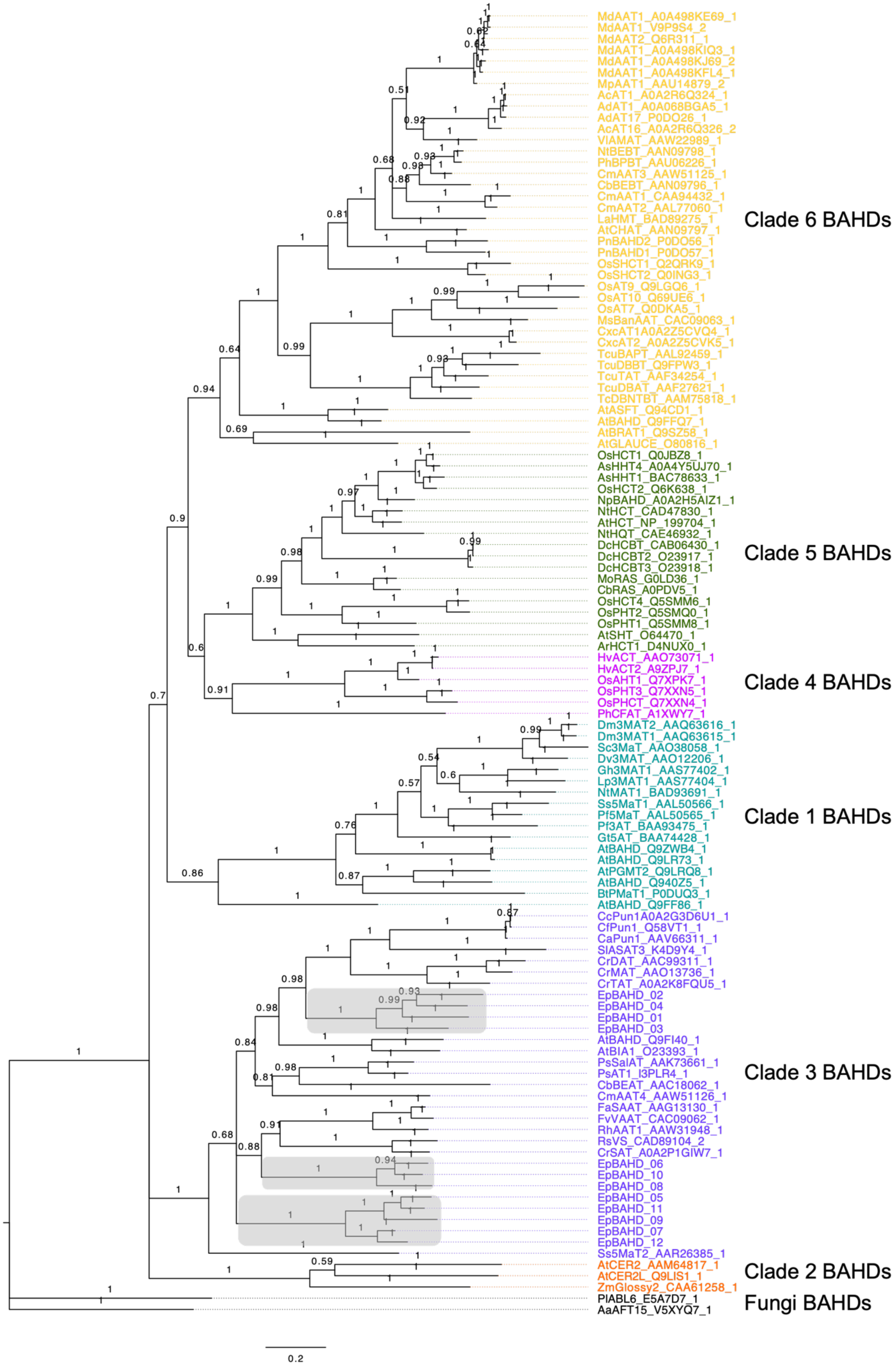
MdAAT1_A0A498KE69_1
MdAAT1_V9P9S4_2
MdAAT2_Q6R311_1
MdAAT1_A0A498KIQ3_1
MdAAT1_A0A498KJ69_2
MdAAT1_A0A498KFL4_1
MpAAT1_AAU14879_2
AcAT1_A0A2R6Q324_1
AdAT1_A0A068BGA5_1
AdAT17_P0DO26_1
AcAT16_A0A2R6Q326_2
VlAMAT_AAW22989_1
NtBEBT_AAN09798_1
PhBPBT_AAU06226_1
CmAAT3_AAW51125_1
CbBEBT_AAN09796_1
CmAAT1_CAA94432_1
CmAAT2_AAL77060_1
LaHMT_BAD89275_1
AtCHAT_AAN09797_1
PnBAHD2_P0DO56_1
PnBAHD1_P0DO57_1
OsSHCT1_Q2QRK9_1
OsSHCT2_Q0ING3_1
OsAT9_Q9LGQ6_1
OsAT10_Q69UE6_1
OsAT7_Q0DKA5_1
MsBanAAT_CAC09063_1
CxcAT1A0A2Z5CVQ4_1
CxcAT2_A0A2Z5CVK5_1
TcuBAPT_AAL92459_1
TcuDBBT_Q9FPW3_1
TcuTAT_AAF34254_1
TcuDBAT_AAF27621_1
TcDBNTBT_AAM75818_1
AtASFT_Q94CD1_1
AtBAHD_Q9FFQ7_1
AtBRAT1_Q9SZ58_1
AtGLAUCE_O80816_1
Clade 6 BAHDs
OsHCT1_Q0JBZ8_1
AsHHT4_A0A4Y5UJ70_1
AsHHT1_BAC78633_1
OsHCT2_Q6K638_1
NpBAHD_A0A2H5AIZ1_1
NtHCT_CAD47830_1
AtHCT_NP_199704_1
NtHQT_CAE46932_1
DcHCBT_CAB06430_1
DcHCBT2_O23917_1
DcHCBT3_O23918_1
MoRAS_G0LD36_1
CbRAS_A0PDV5_1
OsHCT4_Q5SMM6_1
OsPHT2_Q5SMQ0_1
OsPHT1_Q5SMM8_1
AtSHT_O64470_1
ArHCT1_D4NUX0_1
Clade 5 BAHDs
HvACT_AAO73071_1
HvACT2_A9ZPJ7_1
OsAHT1_Q7XPK7_1
OsPHT3_Q7XXN5_1
OsPHCT_Q7XXN4_1
PhCFAT_A1XWY7_1
Clade 4 BAHDs
Dm3MAT2_AAQ63616_1
Dm3MAT1_AAQ63615_1
Sc3MaT_AAO38058_1
Dv3MAT_AAO12206_1
Gh3MAT1_AAS77402_1
Lp3MAT1_AAS77404_1
NtMAT1_BAD93691_1
Ss5MaT1_AAL50566_1
Pf5MaT_AAL50565_1
Pf3AT_BAA93475_1
Gt5AT_BAA74428_1
AtBAHD_Q9ZWB4_1
AtBAHD_Q9LR73_1
AtPGMT2_Q9LRQ8_1
AtBAHD_Q940Z5_1
BtPMaT1_P0DUQ3_1
AtBAHD_Q9FF86_1
Clade 1 BAHDs
CcPun1A0A2G3D6U1_1
CfPun1_Q58VT1_1
CaPun1_AAV66311_1
SlASAT3_K4D9Y4_1
CrDAT_AAC99311_1
CrMAT_AAO13736_1
CrTAT_A0A2K8FQU5_1
EpBAHD_02
EpBAHD_04
EpBAHD_01
EpBAHD_03
AtBAHD_Q9FI40_1
AtBIA1_O23393_1
PsSalAT_AAK73661_1
PsAT1_I3PLR4_1
CbBEAT_AAC18062_1
CmAAT4_AAW51126_1
FaSAAT_AAG13130_1
FvVAAT_CAC09062_1
RhAAT1_AAW31948_1
RsVS_CAD89104_2
CrSAT_A0A2P1GIW7_1
EpBAHD_06
EpBAHD_10
EpBAHD_08
EpBAHD_05
EpBAHD_11
EpBAHD_09
EpBAHD_07
EpBAHD_12
Ss5MaT2_AAR26385_1
Clade 3 BAHDs
AtCER2_AAM64817_1
AtCER2L_Q9LIS1_1
ZmGlossy2_CAA61258_1
Clade 2 BAHDs
PlABL6_E5A7D7_1
AaAFT15_V5XYQ7_1
Fungi BAHDs
0.2

**Fig. S10.** Phylogenetic analysis of acyltransferases identified in this study. Sequence alignment was done using Muscle v3.8.425 (19). The displayed gene tree was then constructed with Bayesian analyses using MrBayes v3.2.7a (20). Posterior probabilities were reported as supporting values for nodes in the trees and scale bar represents substitutions per nucleotide site.

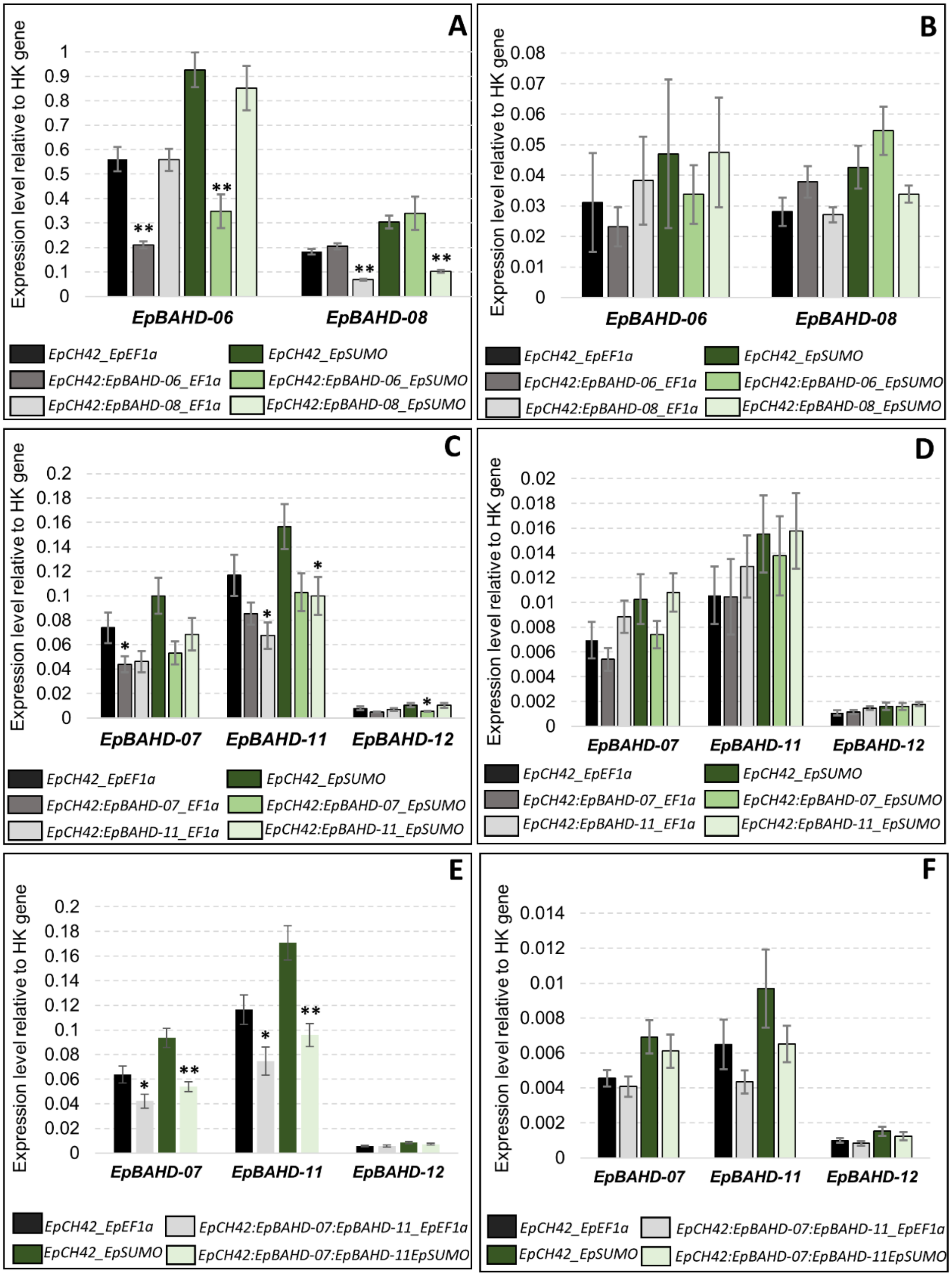


**Fig. S11.** Transcript abundances of selected genes in VIGS experiments. Expression levels of the VIGS-targeted genes and their homologues, relative to two selected housekeeping genes: *EpEF1a* and *EpSUMO* were measured for stem (A, C, E) and leaf (B, D, F) tissues for VIGS marker-only (*EpCH42*, black and dark-green bars) and marker plus BAHD genes (light-grey and light green bars). Error bars – SEM (n = 6 for A,B, E, and F, n = 5 for C and D). Statistically significant (T-test)

changes between relevant control (*EpCH42*) and BAHD silenced genes are indicated by asterisks (**-p-value <0.01, *-p-value <0.05).

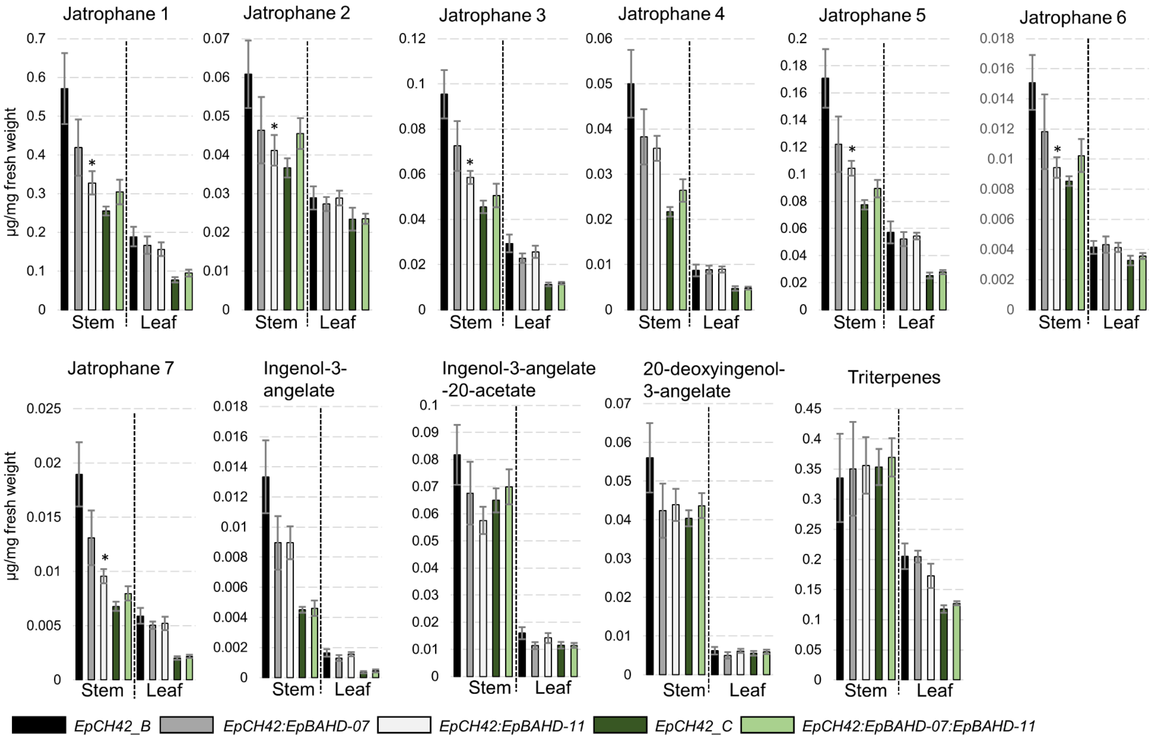


**Fig. S12.** VIGS analysis of *EpBAHD-07* and *EpBAHD-11* in *Euphorbia peplus.* Metabolite levels in VIGS material were measured for stem and leaves in VIGS marker-only (EpCH42_B – black bars and EpCH42_C – dark green bars), marker plus individually silenced BAHD genes: EpCH42 :EpBAHD-07 (dark grey bars), EpCH42:EpBAHD-11 (light grey bars) and marker plus simultaneously silenced BAHD genes EpCH42:EpBAHD-07:EpBAHD-11 (light green bars). Triterpenes represent the sum of four major triterpenes annotated in Datasets S1 and S2. Error bars – SEM (n = 5 for black and grey bars, n = 6 for green bars). Statistically significant (T-test) changes between control (EpCH42_A) and silenced BAHD genes indicated by asterisks separately for each tissue (*- p-value < 0.05; ** - p-value < 0.01).

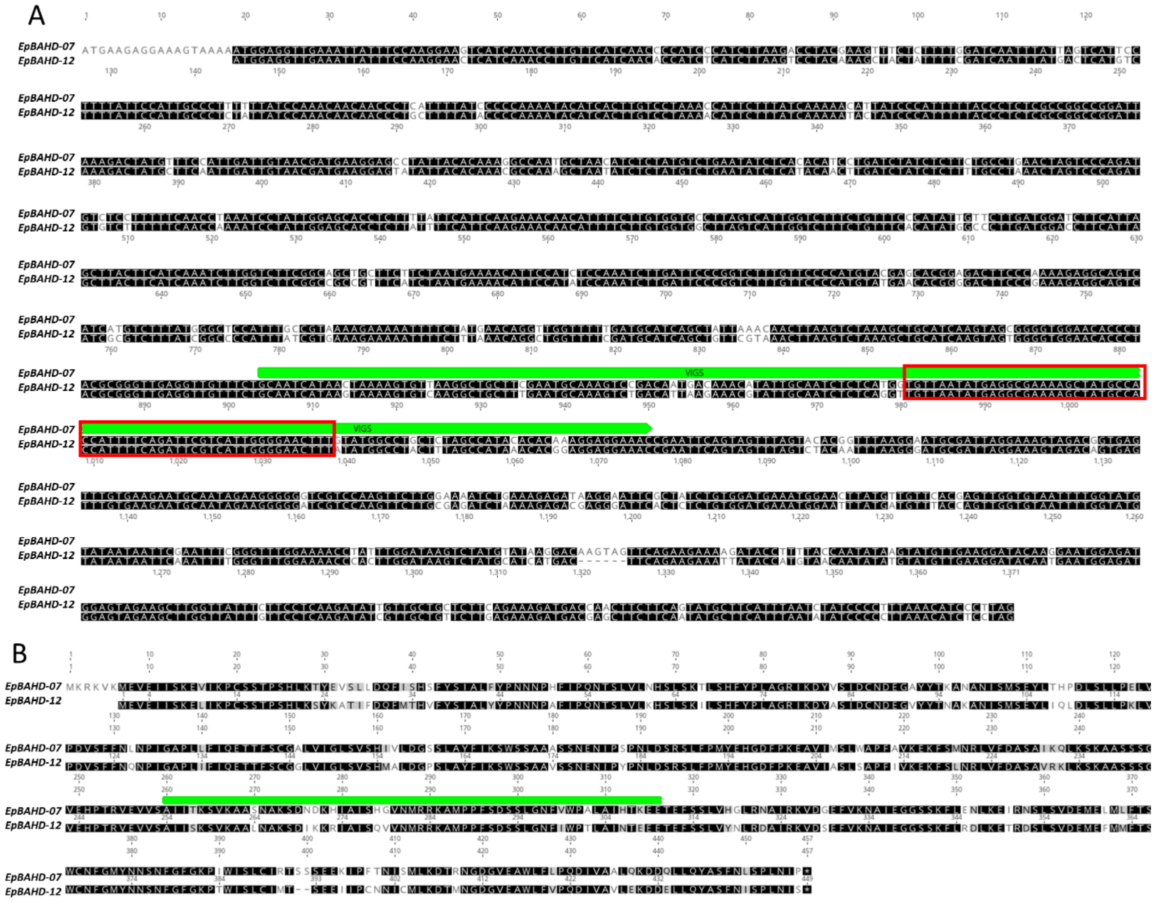

**Fig. S13.** Nucleotide (A) and protein (B) sequence alignment of *EpBAHD-07* and *EpBAHD-12*. Fragment of the *EpBAHD-07* sequence targeted by VIGS is highlighted in green. The 56bp stretch of identical sequence within that VIGS-targeted region is highlighted by the red box (panel A).

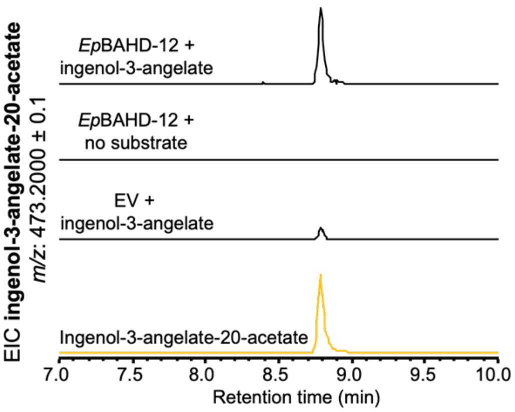


**Fig. S14.** Transient expression of *Ep*BAHD-12 in *N. benthamiana* in the presence of ingenol-3-angelate (**1a**) leads to formation of ingenol-3-angelate-20-acetate (**2**).

**Table S1.** Co-expression analysis of annotated BAHD-acyltransferases with EpCAS. The Pearson correlation coefficients (r) are displayed.

| **Transcript ID** | **Proposed Function** | ***r*** |
|---|---|---|
| CL2478.Contig1_All | EpBAHD-01 | 0.91 |
| CL1599.Contig12_All | EpBAHD-02 | 0.84 |
| CL9095.Contig3_All | EpBAHD-03 | 0.81 |
| Unigene12606_All | EpBAHD-04 | 0.79 |
| CL4712.Contig8_All | EpBAHD-05 | 0.98 |
| CL1163.Contig1_All | EpBAHD-06 | 0.97 |
| CL3102.Contig3_All | EpBAHD-07 | 0.97 |
| CL1163.Contig8_All | EpBAHD-08 | 0.96 |
| Unigene4204_All | EpBAHD-09 | 0.96 |
| Unigene21553_All | EpBAHD-10 | 0.96 |
| CL4712.Contig6_All | EpBAHD-11 | 0.94 |

**Table S2.** Oligonucleotide primers used in this study.

| Target Seq. ID | GenBank Accession | Primers (5´→3´) | Purpose |
|---|---|---|---|
| *Primers used for cloning of acyltransferase gene candidates into 3∧1 for transient expression in N. benthamiana* | | | |
| EpBAHD-01F | PQ801599 | TTTATGAATTTTGCAGCTCGATGCAGCCTAAGGTAAATTCCAAAG | Forward primer for cloning into 3∧1 vector |
| EpBAHD-01R | PQ801599 | GACAACCACAACAAGCACCGTTAAACATAAGTGTCCAAGTC | Reverse primer for cloning into 3∧1 vector |
| EpBAHD-02F | PQ801600 | TTTATGAATTTTGCAGCTCGATGGCCCCAAAATTTCATCCTG | Forward primer for cloning into 3∧1 vector |
| EpBAHD-02R | PQ801600 | GACAACCACAACAAGCACCGTTAAACGCAAGTATCCAAG | Reverse primer for cloning into 3∧1 vector |
| EpBAHD-03F | PQ801601 | TTTATGAATTTTGCAGCTCGATGACCCCAAAAATGCTGCC | Forward primer for cloning into 3∧1 vector |
| EpBAHD-03R | PQ801601 | GACAACCACAACAAGCACCGTTAATCAATCAAACAATCTTCAAG | Reverse primer for cloning into 3∧1 vector |
| EpBAHD-04F | PQ801602 | TTTATGAATTTTGCAGCTCGATGAAGCCTGAGGTAGTTTCAAG | Forward primer for cloning into 3∧1 vector |
| EpBAHD-04R | PQ801602 | GACAACCACAACAAGCACCGTTAAATGCTTGGATTGACCG | Reverse primer for cloning into 3∧1 vector |
| EpBAHD-05F | PQ801603 | TTTATGAATTTTGCAGCTCGATGATGGGATCAAATTGGAAAATCAAAATG | Forward primer for cloning into 3∧1 vector |
| EpBAHD-05R | PQ801603 | GACAACCACAACAAGCACCGTTATAATGGAGATGGATTAAATGAAG | Reverse primer for cloning into 3∧1 vector |
| EpBAHD-06F | PQ801604 | TTTATGAATTTTGCAGCTCGATGCAGATAAAAGTAGAGTTGATATC | Forward primer for cloning into 3∧1 vector |
| EpBAHD-06R | PQ801604 | GACAACCACAACAAGCACCGTCATAAGCGTGAATACAAATTTG | Reverse primer for cloning into 3∧1 vector |
| EpBAHD-07F | PQ801605 | TTTATGAATTTTGCAGCTCGATGGAGGTTGAAATTATTTCCAAGG | Forward primer for cloning into 3∧1 vector |
| EpBAHD-07R | PQ801605 | GACAACCACAACAAGCACCGCTAAGGGATGTTTAAAGGGG | Reverse primer for cloning into 3∧1 vector |
| EpBAHD-08F | PQ801606 | TTTATGAATTTTGCAGCTCGATGGGGATTAAAGTAGAGTTCATTTC | Forward primer for cloning into 3∧1 vector |
| EpBAHD-08R | PQ801606 | GACAACCACAACAAGCACCGTCATAATCGTGAATTCAAGTTG | Reverse primer for cloning into 3∧1 vector |
| EpBAHD-09F | PQ801607 | TTTATGAATTTTGCAGCTCGATGATTCAAGAAACAACATTTTCTTGTG | Forward primer for cloning into 3∧1 vector |
| EpBAHD-09R | PQ801607 | GACAACCACAACAAGCACCGTCAACAAGAACTTAAGGGAG | Reverse primer for cloning into 3∧1 vector |
| EpBAHD-10F | PQ801608 | TTTATGAATTTTGCAGCTCGATGGAGATCCCAGTAGAGTTC | Forward primer for cloning into 3∧1 vector |
| EpBAHD-10R | PQ801608 | GACAACCACAACAAGCACCGTCAAGTTTGGATTGTATTTGG | Reverse primer for cloning into 3∧1 vector |
| EpBAHD-11F | PQ801609 | TTTATGAATTTTGCAGCTCGATGGGATCAAACAGGAAAATTAAAATGG | Forward primer for cloning into 3∧1 vector |
| EpBAHD-11R | PQ801609 | GACAACCACAACAAGCACCGTTAAATACTTAAAGGAGATGG | Reverse primer for cloning into 3∧1 vector |
| EpBAHD-12F | PQ801610 | TTTATGAATTTTGCAGCTCGATGGAGGTTGAAATTATTTCCAAGG | Forward primer for cloning into 3∧1 vector |

| | | | |
|---|---|---|---|
| EpBAHD-12R | PQ801610 | GACAACCACAACAAGCACCGCTAG GAGATGTTTAAGGGGG | Reverse primer for cloning into 3∧1 vector |
| *Primers used for cloning of acyltransferase gene candidates into popinF for recombinant protein production* | | | |
| EpBAHD-06F | | AAGTTCTGTTTCAGGGCCCGATGC AGATAAAAGTAGAGTTGATATCC | Forward primer for cloning into pOPINF vector |
| EpBAHD-06R | | ATGGTCTAGAAAGCTTTATCATAAG CGTGAATACAAATTTGC | Reverse primer for cloning into pOPINF vector |
| EpBAHD-08F | | AAGTTCTGTTTCAGGGCCCGATGG GGATTAAAGTAGAGTTCATTTC | Forward primer for cloning into pOPINF vector |
| EpBAHD-08R | | ATGGTCTAGAAAGCTTTATCATAAT CGTGAATTCAAGTTGGG | Reverse primer for cloning into pOPINF vector |
| EpBAHD-07F | | AAGTTCTGTTTCAGGGCCCGATGG AGGTTGAAATTATTTCCAAGG | Forward primer for cloning into pOPINF vector |
| EpBAHD-07R | | ATGGTCTAGAAAGCTTTACTAAGG GATGTTTAAAGGGGATAG | Reverse primer for cloning into pOPINF vector |
| EpBAHD-11F | | AAGTTCTGTTTCAGGGCCCGATGG GATCAAACAGGAAAATTAAAATG | Forward primer for cloning into pOPINF vector |
| EpBAHD-11R | | ATGGTCTAGAAAGCTTTATTAAATA CTTAAAGGAGATGGATTAATTG | Reverse primer for cloning into pOPINF vector |
| *Primers used for cloning and creating VIGS construcs and for qPCR* | | | |
| EpBAHD-06_Vigs_F | | CGGTACCGAGCTCACGCGTCGCTT TCTTATTTAGTCGGATAGTG | Forward primer with XhoI tail for cloning into pTRV2 vector |
| EpBAHD-06_Vigs_R | | TTTAATGTCTTCGGGACATGCCCA CTCGTATTATCAATCAATGTCGT | Reverse primer with SmaI tail for cloning into pTRV2 vector |
| EpBAHD-07_Vigs_F | | CGGTACCGAGCTCACGCGTCGCA ATCATAACTAAAAGTGTTAAGGCT | Forward primer with XhoI tail for cloning into pTRV2 vector |
| EpBAHD-07_Vigs_R | | TTTAATGTCTTCGGGACATGCCCG TTTCCTCCTTTGTGTGTATGG | Reverse primer with SmaI tail for cloning into pTRV2 vector |
| EpBAHD-08_Vigs_F | | CGGTACCGAGCTCACGCGTCGTTT TCATATTAAGTCGGGTAAGAG | Forward primer with XhoI tail for cloning into pTRV2 vector |
| EpBAHD-08_Vigs_R | | TTTAATGTCTTCGGGACATGCCCAT CCATATTATCAATCACTGTTACAC | Reverse primer with SmaI tail for cloning into pTRV2 vector |
| EpBAHD-11_Vigs_F | | CGGTACCGAGCTCACGCGTCCTG CTCTGAACACAAAGTCA | Forward primer with XhoI tail for cloning into pTRV2 vector |
| EpBAHD-11_Vigs_R | | TTTAATGTCTTCGGGACATGCCCT CTGTGGAGAACGCGTG | Reverse primer with SmaI tail for cloning into pTRV2 vector |
| qRT-EpBAHD-06_F | | AGCTTGACAGAGGACGAC | Primers for qPCR amplification of the genes targeted by VIGS |
| qRT-EpBAHD-06_R | | TCATAAGCGTGAATACAAATTTGC | |
| qRT-EpBAHD-07_F | | GACAAGTAGTTCAGAAGAAAAGAT AC | |
| qRT-EpBAHD-07_R | | CAGCAACAATATCTTGAGGAAG | |
| qRT-EpBAHD-08_F | | GTTAGCTTGACAGAGAAAGAAATG | |

| | | | |
|---|---|---|---|
| qRT-EpBAHD-08_R | | TCGTGAATTCAAGTTGGGATTC | |
| qRT-EpBAHD-11_F | | GAAGATGGTGAAGAGATGCC | |
| qRT-EpBAHD-11_R | | CTTAAAGGAGATGGATTAATTGAAGC | |
| qRT-EpEF1a_F | OL744076 | CTGGCATAATTAAGATGATTCCG | Primers for qPCR amplification of housekeeping genes |
| qRT-EpEF1a_R | OL744076 | CTTCTTCTTGGCAGCAGATT | |
| qRT-EpSUMO_F | OL744078 | CCAGGACCTGCTCGATC | |
| qRT-EpSUMO_R | OL744078 | GATATAATCGTTTTCACATCAGACC | |
| qRT-EpBAHD-12_F | | TGGATAAGTCTATGCATCATGACTTC | Primers for qPCR amplification of the genes closely related to *BAHD's* sequences targeted by VIGS. |
| qRT-EpBAHD-12_R | | GGATATATTAAATGAAGCATATTGAAGAAG | |

**Table S3.** Amino acid sequences of acyltransferase genes used in this study.

| Acyltransferase | Amino acid sequence |
|---|---|
| *Ep*BAHD-01 | MQPKVNSKEFVKPSPSNHPKIHYLSFFDQMVNLNSYICLLFFYTSKNRQTSAHNNTSSVLKTSLSAALSRFYPLAGRLK<br>GGLTMDCNDEGALFLESKLECSLSELLNNPDLEMLKPLFPDNLPFNFSSNIPLAVQVTYFQCGGMSIGVCVSHKILDM<br>RGMSIFIKEWSSLARHPEKEIHLEFNIGSFYPPLDLPILQEKVEPPSEMEDCVHRRVVFDGSKIAKLKEMVVDQVPNPT<br>RVEVVISLLYRSALYANAKATSSCSIVPSFMTHAMSLHTKVSPPLSERSRGNIWGMFIVVPGQDERDIELRYLVKEFRN<br>AKTEMSNSCANNNKEDLCQMALKNMKYLSTYANEYQVYTCSSWCRFPFYESDFGWGKPLWVTTTCMKLKNFIFLED<br>TREGDGIEALISLEGKAMAIFEKDQELLSFCRFNPSVLDLDTYV |
| *Ep*BAHD-02 | MAPKFHPEIVSTEIVKPSPNNHPSFHTLSLFDQLNHPVYVPFLFFYTSNKNGDEFHKSNVLKTSLSDALLHYYPLAGRIK<br>DDVTVDCNDEGTLFLEARIACGLSELLKNHDPDTLKVLLPDGVSGKDSTSKSPVVVQVTYFQCGGMSIGLCLCHKILD<br>MASMTYFIYDWASLARKSGDEIRPEFNAGSLYPPLDLPILKQKQAVFEDMKDCVDRRFVFYGPKIDKLKEMVADQVEK<br>PTRVEVVASLLYKSAISANAKARKSGSIEPSVLYLPMNLRNKVAQLSERHRGNILGTFKVPVRDEREIELGLLVKQFRK<br>VKAEYADLCASKTNSEDLCPFITELLIDGSLENLNVDHDVYSLTSFCRYPFYGADFGWGKPLWVTFPNIKMKNVMFCL<br>DTKEGDGIDVCIVLEEKAMSIFENDQELLSFCSINPSVLDTCV |
| *Ep*BAHD-03 | MTPKMLPEIVSREIIKPPSTTQNNHPTTHNLSFFDQTSPCVYVPLILFYSSNNSETNHHKSGVLKASLSATLSEFYPLSG<br>RTKDDVTVDCNDEGVVFLEANVGCNLSEMLKCPDDETLKLLLPDGLYYKDFRLSSPVVVQITYFESGGMSIGLCLSHK<br>VQDMESICAFLKKWASIARKSDQDIAPEFNIAARYPPLDLPFLKEQQPAQEMKNCASRRVVFRSESIAKLKEIAGDHVQ<br>NPTRVEVVASLLYKSATCAVSKTSSSGNTLPRTSMHHVMNLRTKVSPPLSDRKAGNLTGVFPVSAPMGTEFGELVKE<br>MRQEKMWFSDSCTKISNKEELYTFVLESMKGLGTYFSDENEEIYFCSSWCRYPFYEIDFGWGKPFWTTTVFWEMKN<br>LILMIDDQGGDGIEAFVTLDKKAMDLFETDAEILEFSRINPCVDLEDCLID |
| *Ep*BAHD-04 | MKPEVVSREIVKPSSSIPKNHPKFHNLSFFDQIVPSYYVPLLYFYTCKNEEPDHNLRSRVLKTSLSAALSHYYPLAGRIN<br>YDMVSVDCNDEGALFLEATIGCGLSEILISPDDVILKLLFPDGLCYKDSTQTGPVAVQVTYFQCGGMSVGICLCHKMME<br>MASMSSFISDWASLARNSDKEIDPEFNIGSLYPPLDLPILKQKQASPPVTKDCVSRRVVFDGLKIAKLKELVAERVPNPT<br>RVEVVASLLYKSAICGNAKASSTGSLIPSFINCAMNLRSRVSPPISEREKGNISGIFRVSVRDEREVELGCLVTEFRKAK<br>TRLSNSCANITSTEDLSQLILKTMKPLSEYFNAEHEVYGCTSWCRYPFYGTDFGWGKPLWVTTVLYKLKNGMTFIDTK<br>DGGGIEAFICLEDKAVTIFENDQDLLLFCSVNPSI |
| *Ep*BAHD-05 | MMGSNWKIKMEVEIISRGCIKPSSSTPSHLKSYKISLLDQFMPPIYYSIALFYPNSNNDLTISQRSRLLKHSLSKTLSHFY<br>PLAGKIKDYLSVDCNDEGAYYIDAKANIPLSEFLTRPDHTALRKFVPDTSPFNANPIGAYVVMFQETTFSCGGLALGIAA<br>SHMVLDGISLASFIKAWASSTSASEKEIPYPNLDSPSVFPHIEDFPGDALFPALWIPFVKTGKLAIRRLMFDATSIDKLKA<br>KASPSVVNPTRVEVVSAIITKTIKAALNAKSIAFVHGVSMRRKATTPFPDCSLGNFIWNVHAYSGENESQISSLAYNLRN<br>VIRKVDNEFVNNIATKGFSGLYEEIKEIKKGLSSDGMELISFSSWCNFGVYDNSDFGFGKPIWMSYCTISDGEEMPFFN<br>MGALNDTRSGNGVEVWLYLSEDVAAFVENDDELLQYASFNPSPL |
| *Ep*BAHD-06 | MQIKVELISKELIKPSSPTPLHLHKLEFSFIDQNQYPLSFPFVLFYEKSNIPNQQRSNILKKSLSQALTIFYPLAGRINNYS<br>YADCNDEGALFIEASANCQLSDILRNRNQYHNCNRNFIPVQPDKGVHQYGSLFQITYFSCGGLAVAFAMPHMLGDAL<br>SQFIFLNGWAAVARGIKVDVPPIGSASIFPPRNIPGFDLSQWVFKEKVITKPFVFDALTLSALKEKYSANGEKFSRVQAL<br>CAFLFSRIVASISQAKDARYMAIYSMNVRQILDPPVPKQSFGNLVWAATTLIDNTSKEDEIARKIKDSVKGVNAESLNKL<br>QNGELSFSKEAFMDSLKGEIVRYSFTSLCNFPTYDVDFGWGKPEWVTACGLVIDNLVICVDTKGRGIEAWISLTEDDM<br>TKFEKDKVLVSHLSSNANANLYSRL |
| *Ep*BAHD-07 | MEVEIISKEVIKPCSSTPSHLKTYEVSLLDQFISHSFYSIALFYPNNNPHFIPQNTSLVLNHSLSKTLSHFYPLAGRIKDYV<br>SIDCNDEGAYYTKANANISMSEYLTHPDLSLLPELVPDVSFFNLNPIGAPLLFIQETTFSCGALVIGLSVSHIVLDGSSLA<br>YFIKSWSSAAASSNENIPSPNLDSRSLFPMYEHGDFPKEAVIMSLWAPFAVKEKFSMNRLVFDASAIKQLKSKAASSS<br>GVEHPTRVEVVSAIITKSVKAASNAKSDNDKHIAISHGVNMRRKAMPPFSDSSLGNFVWPALAIHTKEETEFSSLVHGL<br>RNAIRKVDGEFVKNAIEGGSSKFLENLKEIRNSLSVDEMELMLFTSWCNFGMYNNSNFGFGKPIWISLCIRTSSSEEKI<br>PFTNISMLKDTRNGDGVEAWLFLPQDIVAALQKDDQLLQYASFNLSPLNIP |
| *Ep*BAHD-08 | MGIKVEFISKKTIKPSSPTPHHLRKLKFSFFDQIQFPISIPFIFFYEKSNISNEERSNLLKQSLSKALTIFYPLAGRINNHSYA<br>DCNDEGGLFIEAKANCKLSDILGDRNQYHNNSEKFIALQPDKGIHQVGSLFQITSFTCGGLAVSFAMPHMLGDAVSIFA<br>FMNGWAAIARGDTIDVPSFNAASIFPPKSSSGIDFSQMVFKDNVVTKSFVFEASAISAIRDKYSANGEKLSRAQTLYVFI<br>LSRVRASTSRAKGGNNNRYIVVNSVNVRQMLDLPVSKQSFGNFIWAGVTVIDNMDLIEKENGCHELGKRIKDSIKSVN<br>AELLRKLKNGEFSSNIEALVESIEGEIIRFSLTSICNFPVYEVDFGWGKPEWVATSALFFDNFVVLLDTKDGQGIEAWVS<br>LTEKEMAMFENDKMLISHTSSTPNRNPNLNSRL |

| | |
|---|---|
| *Ep*BAHD-09 | MIQETTFSCGGLALGFSVSHMVLDGTSFASFIKAWASSASGNKIPYPNLDSSSVFPQFEAFPNDAICSSLWTPFARKG<br>KLAVRRLMFDQSSIDRLKVKASTSVKSPTRVEVVTSIITKRVKAALNVKSVAISHSVNIRRKATTPDYSLGNIVLMVHAFF<br>TDDKESEIRILRNAIRKVDNDFVKIKAIEGGFSELYQEHKEMSSRLSINEMELISFTSWCNFGLYNDSDFGWGKPVWIS<br>HNSLFEGEVMPFYNMCILNDTRSGNGVEAWLYSSQDVADFIGKDDELLQYASINPSPLSSC |
| *Ep*BAHD-10 | MEIPVEFITKELIKPSSPTPPHLRELKFSFIDQNQYPVSIPFVFFYEKSTIPNHERYNLLKQSLSQALTIFYPLAGRINNYD<br>YADCNDEGALIVQTKAHCQLSDILENRNQYHYCYKKFIPLQPHKGVHQYGSLFQITYFNCGGLAISFALPHMLGDALSQ<br>FIFLNGWAAVARGNQVDIPPFVSDSIFPPRNIPGFDLSQWVFKENIVTKSFVFDAFAISALRDKYSTGGETISRVQALCV<br>FLFTRIIASLSRQAKDNRFMAIHSVNVRRKLDPPVPDQSFGNLVWAATSVIDNISEEDEIARRIKDSIKGVNAEPLNKLQK<br>GELSFSKETFMESLKGEIVRYSFSSLSNFPVYDVDFGWGKPEWVTTSGVIIDNLVMCIDTKEGRGIEAWVNLTEEDMA<br>KFEKDKVLLSHLSSAPNTIQT |
| *Ep*BAHD-11 | MGSNRKIKMELEIISKECIKPSSSTPSHLQTYKISLLDQLMLPRYYSIALFYPNSNNDLTLTQRSRLLKHSLSKTLSHFYP<br>LAGKIKDYLSIDCNDEGAYYIEAKVNIPLSEFLTRPDHTSLKKFVPESSLFNANPIGAYIVMIQETTFSCGGLALGIVVSHM<br>VLDAFSFASFVKAWASLASDKQIPYPNLDSPSVFPQLEGFSANTLFPALWGPFAKKGKLAMRRLVFDASGIDKLKAKA<br>SPSVENPSRVEVVSAIITKTIKAALNTKSIAFTQAVNMRRKAMTPFPDCSLGNFIWSVHAFSTENEDQISSLVHNLRNAV<br>RKVDNKFVNNIATEGGISKLYEEMEEIKKRLSSDGMELVNFSSWCNFGVYDNTDFGFGKPLWMIYCIVEDGEEMPFYS<br>MCNMNDTRSGNGVEAWLYLSQDVAAFVENDVELLQYASINPSPLSI |
| *Ep*BAHD-12 | MEVEIISKELIKPCSSTPSHLKSYKATIFDQFMTHVFYSIALYYPNNNPAFIPQNTSLVLKHSLSKILSHFYPLAGRIKDYA<br>SIDCNDEGVYYTNAKANISMSEYLIQLDLSLLPKLVPDVSFFNQNPIGAPLIFIQETTFSCGGLVIGLSVSHMALDGPSLA<br>YFIKSWSSAAVSSNENIPYPNLDSRSLFPMYEHGDFPKEAVIASLSAPFIVKEKFSLNRLVFDASAVRKLKSKAASSSG<br>VEHPTRVEVVSAIISKSVKAALNAKSDIKKRIAISQVVNMRRKAMPPFSDSSLGNFIWPTLAINTEEETEFSSLVYNLRDA<br>IRKVDSEFVKNAIEGGSSKFLRDLKETRDSLSVDEMEFMMFTSWCNFGMYNNSNFGFGKPTWISLCIMTSEEIIPCNNI<br>CMLKDTMNGDGVEAWLFVPQDIVAVLEKDDELLQYASFNISPLNIS |

**Dataset S1 (separate file).** Levels of selected di- and triterpene metabolites in marker-only (EpCH42) and marker plus EpBAHD-06 and EpBAHD-08 silenced (EpCH42:EpBAHD-06 and EpCH42:EpBAHD-08) stems and leaves.

**Dataset S2 (separate file).** Levels of selected di- and triterpene metabolites in marker-only (*EpCH42*) and marker plus *EpBAHD-07* and *EpBAHD-11* silenced (*EpCH42:EpBAHD-07and EpCH42:EpBAHD-11*) stems and leaves.

**Dataset S3 (separate file).** Levels of selected di- and triterpene metabolites in marker-only (*EpCH42*) and marker plus *EpBAHD-07* and *EpBAHD-11* silenced (*EpCH42:EpBAHD-07and EpCH42:EpBAHD-11*) stems and leaves.
